# ACCUMULATION OF MICROPLASTIC AND MICRORUBBER PARTICLES IN STORMWATER POND FISH AND INVERTEBRATES

**DOI:** 10.1101/2022.03.03.482888

**Authors:** Stephanie B. LaPlaca, Peter van den Hurk

## Abstract

Stormwater ponds serve as best management practices to trap or remove both physical and chemical pollutants before further discharge of stormwater into receiving natural waterbodies. Stormwater ponds can therefore accumulate high levels of pollutants, including microplastics (MP) and microrubber (MR) from road runoff. It was hypothesized that biota in stormwater ponds that receive large amounts of road runoff, or are in proximity to roadways and major developments, would contain significant amounts of MP per individual, with high abundances of tire particles (TP). Fish and invertebrates were collected from five stormwater ponds and their adjacent tidal creeks in Mt. Pleasant, SC, USA. Whole organisms were digested using KOH and digested contents were filtered and analyzed by visual microscopy to identify and count MPs.

The majority (>80%) of MP recovered from biota across all sites were suspected TP. The average number of MP per individual ranged from 0.3 to 71 MP and the average number of suspected tire particles per individual ranged from 0 to 57.7 tire particles.

There were significant differences in MP per individual observed between sites and between species. A combination of factors such as availability of MPs based on surrounding land use, pond hydrodynamics, organism size, and species feeding habitat likely influenced the total MP observed among the different sites and species analyzed. These data provide preliminary examination into the fate and transport of MP and MR in stormwater ponds and an evaluation of MP and MR abundance in organisms from stormwater ponds in the coastal zone.

## Introduction

Recent publications have drawn attention to the prevalence of microplastic (MP) particles in air, sediment, and water samples worldwide (Eerkes-Medrano et al., 2015; Gasperi et al., 2018). A variety of studies cite MPs in field-collected organisms, including terrestrial and aquatic species (Foekema et al., 2013; He et al., 2013). More recently, microrubber (MR) has been identified as a secondary microplastic type of concern, entering the environment from tire wear under typical driving conditions. Studies have sampled MR from air, sediment, and water as well; however, fewer data on the abundance of MR particles in biota are available (Wik &Dave, 2009). Microplastic and microrubber particles are assumed to coincide in environmental samples and as synthetic polymer particles have similar transport mechanisms. Therefore, it is assumed that MR particles are as ubiquitous in biota as MP. As a variety of studies has demonstrated that MR can have serious toxicological effects on aquatic organisms, understanding the presence and abundance of MR in field-collected organisms is essential for characterizing the overall environmental risk of MR in the environment (Cunningham et al., 2021; Halle et al., 2020; LaPlaca &van den Hurk, 2020; Panko et al., 2013).

Land-based activities provide significant contributions to MPs in the aquatic environment with sources from industrial, commercial, and residential areas. More specifically, sources of MPs include textile washing, personal care and cosmetic products, tires, artificial turf, road paints, packaging, littering, and the construction industry (Xu et al., 2020). In addition to transporting MPs, stormwater runoff is widely recognized as a major transporter of contaminants such as metals, organic pollutants, nutrients, and pathogens from the surrounding landscape to receiving waters (Hoffman et al., 1984; US EPA, 2012). For many pollutants including MPs, stormwater retention ponds serve as a hot spot for pollution and play a role in transport of MPs from land to the aquatic environment (Liu et al., 2019).

Almost 40% of the total U.S. population lives in coastal counties, yet the coastal zone accounts for less than 10% of the nation’s land mass (NOAA, 2018). South Carolina has experienced rapid coastal population growth rates over the last forty years, resulting in an extraordinary increase in land development along its coast. From 1973 to 1994 in Charleston alone, the population increased by 40% while urban area increased by 240%, and that number is likely even greater at present (Charleston Waterkeeper, 2014). Rapid urbanization of coastal regions presents stormwater management challenges as impervious surfaces dramatically increase in the expanding urbanized area. Stormwater ponds are the most widely used best management practice for mitigating stormwater flooding and pollution control.

The Charleston Harbor is a major urban estuary located on the southeastern Atlantic coast of the U.S., with the City of Charleston residing on the peninsula between the Ashley River and the Cooper River (Crist et al., 2019). The Charleston metropolitan area has a strong economy due to the Port of Charleston, tourism, military installations, medical facilities, and manufacturing. The port is ranked 6^th^ in the U.S. in terms of cargo value (S.C. Ports Authority, 2020). According to the U.S. Census Bureau, the population in the Charleston area (including Goose Creek, Hanahan, North Charleston, Mount Pleasant, and Charleston) is approximately 422,000 individuals (U.S. Census Bureau, 2020).

Utilizing satellite imagery, an estimated 21,594 ponds have been counted within the coastal zone of South Carolina associated with either rural or development-related land uses (Smith et al., 2020). Of those, 43% were determined to be associated with development including residential, golf course, or commercial uses. The Charleston area has approximately 4,000 ponds alone as of 2015, with nearly half of those associated with developments listed above (Smith et al., 2020). In the coastal region of South Carolina, wet detention ponds are the most common type of stormwater pond constructed (Crawford et al., 2010; Drescher et al., 2007). Stormwater ponds remove pollutants through physical, chemical, and biological processes such as sedimentation, photodegradation, or particulate trapping by vegetation, respectively (Beckingham et al., 2019). The densities of MR particles in stormwater runoff, including tire wear, crumb rubber, bitumen particles, range from 1.13 – 2.2 g/cm^3^ and are likely to settle and accumulate in stormwater pond sediments; however, pond hydrodynamics (i.e., size, depth, shape, flow, hydraulic residence time) may influence their settlement (Degaffe &Turner, 2011; Kayhanian et al., 2012; Kell, 2020; Rhodes et al., 2012; Unice et al., 2019).

Another factor related to pollutant and MR accumulation in stormwater ponds is the efficiency of particle capture on the surrounding landscape (e.g., by rain gardens), by stormsewer catch basins, and manufactured treatment devices (MTDs) (Werbowski et al., 2021). MTDs function as stormwater treatment devices before the stormwater is discharged off-site or to receiving water bodies through processes as filtration, screening, or settling (SC DOT, 2015). MR particles that are not effectively captured and sequestered along stormwater runoff pathways may remain suspended and are transported to adjacent receiving waters, like marshes or tidal creeks found in the local area of the current study.

Settled MR particles may be ingested by benthic or bottom-feeding organisms in stormwater ponds or adjacent tidal creeks (Redondo-Hasselerharm et al., 2018). Parker et al. (2020) observed MPs and suspected tire wear particles in the digestive tract of pelagic fish collected from an urban harbor, which suggests that suspension or resuspension of MR in stormwater ponds could also lead to uptake by a variety of species. Feeding ecology is suspected to have an impact on MP exposure and ingestion in aquatic organisms, based on food preference, habitat selection, feeding mechanism and selectivity, although there have been mixed results on the impact of feeding traits on microplastic exposure among various species (Parker et al., 2020; Pazos et al., 2017; Walkinshaw et al., 2020).

In an assessment of MPs in coastal South Carolina (Charleston Harbor, SC, USA), black fragments classified as tire wear particles (TWP) comprised the majority of total microplastics encountered in sediment samples (Gray et al., 2018). It is likely that hot spots of high MR levels occur in proximity to roads and bridges when considering the fate and transport of MR. Evaluation of data from previous studies does indeed indicate high abundances of MR in proximity to roadways, (Järlskog et al., 2020). Additionally, stormwater ponds which function to collect stormwater runoff from roads and other impervious surfaces are also a potential hot spot for MR particles. Kell (2020) measured levels of MP and MR in stormwater ponds in the Charleston area of SC and found that levels decline in pond sediments from stormwater entrance to discharge point at the tidal creek, and then downstream or upstream in the receiving tidal creek, demonstrating that stormwater ponds are an important sink for MR particles within stormwater transport pathways, yet are not perfectly efficient at their capture.

The objective of the present study was to characterize the abundance of MPs found in aquatic biota from stormwater ponds and their adjacent tidal creeks, with an emphasis on MR particulates or tire particles (TP). These data will add to the growing body of knowledge on the efficiency of stormwater ponds in capturing and retaining MR particulates and provide an assessment of MP, and more specifically MR, in stormwater pond biota.

## Methods

### Sampling sites

Stormwater pond sites and adjacent tidal creeks were selected around Mount Pleasant, South Carolina, USA, within the Charleston Harbor watershed. Ponds selected represented a variety of drainage area land uses, including residential, commercial, highway, and golf course areas. Detailed descriptions of pond locations and receiving waterbodies are provided in Supplemental Information Table S1 and Figures S1-S6. Five stormwater ponds (Whipple Road, Oak Marsh, Oyster Point, Tides Condos, Patriots Point) and their adjacent tidal creeks were targeted to sample for aquatic biota. The reference pond (Patriots Point) is located on a golf course and was expected to have little to no influence from vehicular traffic that would create MR particles, aside from golf carts.

### Biota collection

Aquatic biota were captured from stormwater ponds and adjacent tidal creeks using a seine net (2m length, 5mm mesh). Nets were pulled through the ponds as depth allowed in proximity to the shoreline to collect sufficient organisms, with the goal of at least 5 individuals of the most abundant species per location. Nets were pulled through tidal creeks moving from downstream to upstream (near pond) 1-3 times as needed to collect sufficient organisms. Biota (n = 285) were sampled on April 27^th^ and April 28^th^, 2021 from each of the five stormwater ponds and tidal creeks (see table Table S2 for all species information). However, organisms could not be collected from Oak Marsh pond, Whipple Road pond, or the Tides Condos tidal creek as conditions did not allow for collection using the methods described (limited access, too much debris, etc.) Water quality parameters (salinity, pH, temperature, dissolved oxygen) were recorded at each pond and tidal creek site. Collected fish were euthanized using MS-222 solution (1g/L), separated by species, wrapped in labelled aluminum foil, and placed on ice in the field. Invertebrates (i.e. crawdads and grass shrimp) were separated by species, wrapped in labelled aluminum foil, and placed on ice in the field. Immediately upon returning to the lab, all samples were transferred to a -20°C freezer for storage until further analysis.

### Digestion of samples and isolation of particles

Biota were thawed, weighed (wet weight), and length measured individually. Due to the small size (< 100 mm) of most species, organisms were grouped by species into pools of 3 to 12 individuals for digestion. Whole organisms were placed in a glass beaker with 1 M KOH solution at approximately three times the volume of the organic biological material (Foekema et al., 2013; Lusher et al., 2017; Parker et al., 2020). Beakers were sealed with aluminum foil and digested for a period of three to five days, or until interfering tissue had been fully dissolved, with heat applied for a portion of the digestion period (24-hr at 60°C in a water bath). Following digestion, samples were sieved through a series of stacked metallic brass sieves (500µm and 53µm). Contents from each sieve were washed into separate glass beakers with DI water and covered with aluminum foil until further processing.

### Microplastic and microrubber quantification

Sieved digested samples were washed onto a mixed cellulose ester membrane (Whatman, sterile mixed cellulose ester membranes, color: green with black grid, size:

0.45 µm pore) over a glass vacuum filtration funnel to remove liquids. If samples contained a large amount of digested material, more than one filter was utilized and microplastic counts from all filters of the same sample and sieve size were totaled.

Microplastics were identified and counted under a dissecting microscope (20x-40x, Meiji EMT Tokyo, Japan). Microplastics were classified by type as fibers, suspected tire wear particles or fragments, and colors were noted. Other microplastics (foams, films, spheres) were noted, if observed. Microplastics were counted by reading the filter in a crystallizing dish in a serpentine pattern and analyzing each square on the gridded filter. Microplastics were identified using established criteria for microplastic morphology and confirmed using either the hot needle test or manipulation with a probe (Barrows et al., 2017; Lusher et al., 2017). For microplastics not including suspected tire wear particles, items were considered microplastic under the following criteria: homogeneously colored, no visible cellular or organic structure, equal thickness throughout for fibers, and does not crumble when manipulated with forceps. The hot needle test was used to help distinguish between microplastic pieces and organic matter as needed where, plastic pieces would melt or curl in the presence of a very hot needle and biological or non-plastic materials would not (Barrows et al., 2017; De Witte et al., 2014). Suspected tire wear particles are more difficult to identify as they do not react to a hot needle test and have more variability in appearance and behavior depending on tire age, particle age, and particle structural integrity. In general, particles were classified as suspected tire wear particles under the following criteria: darkly colored (black), elongated, cylindrical, or irregular in shape (cuboidal/angular), partially or entirely covered with road dust (glittery sheen), rough surface texture, rubbery flexibility when manipulated with forceps (Kell, 2020; Kreider et al., 2010; Leads &Weinstein, 2019; Parker et al., 2020). Suspected tire wear particles that disintegrated or turned powdery when probed with a dissecting needle were not counted as part of the total suspected tire particles (TP).

### Quality Assurance/Quality Control

To reduce potential contamination from MP, a number of measures were taken. After thawing field-collected samples, organisms were rinsed with deionized water after weighing and measuring to reduce contamination from any MPs potentially adhered to the outer surface of the organisms prior to digestion. In the laboratory, only stainless-steel or glass equipment was used, except for LDPE wash bottles and nitrile gloves. Samples were kept covered to help eliminate airborne contamination. Additionally, all equipment (i.e., beakers, sieves) was rinsed with deionized water three times before use and in between each sample.

Procedural blanks (without tissue) were processed with each batch of samples (n=10 total blanks). Briefly, DI water was poured through the rinsed nested sieves similarly to sample processing, and DI water rinse was collected from the sieve in a glass beaker (approximately 200 mL). Blanks were poured over a gridded filter under a vacuum as described above. The number of microplastics on the gridded filter was enumerated for each blank by visual microscopy.

### Statistical analysis

All statistical analyses were performed using JMP Pro 14 (SAS Institute Inc.) statistical software. Microplastic counts were normalized to the number of individuals examined in each batch processed. The percentage of microplastic types (fiber, suspected tire particle, and fragment) was determined for each site sampled. Data was log transformed to better approximate for normality when required. When the data was approximately normally distributed, an ANOVA or t-test was performed to test for significant differences. Tukey’s post hoc analysis was performed if significant differences were found. Simple linear regression was used to assess relationships between variables, such as MP abundance and biota weight. A p-value of < 0.05 was considered significant.

## Results

### Background Contamination

Procedural blanks (n=10) contained an average of 1.1 ± 0.6 MP per blank. MP sample counts were adjusted to account for procedural blank contamination by subtracting the average counts in procedural blanks from the total MPs counted for each batch analyzed.

### Microplastic counts

A total of 285 organisms were processed for total microplastics which included nine species of fish, one species of grass shrimp, and one species of crawdad (Table S2). Overall, 100% of the samples examined contained individuals with at least 1 MP, with an average of 9.5 ± 6.5 MP per individual across all organisms and all sites sampled (**Figure 1**). As individuals were grouped into batches of 3-12 for processing, the exact number of MP per each individual analyzed, or the percentage of individuals examined overall that contained MP, could not be deduced. A linear regression between microplastics per individual and the average length (mm) and weight (g) of biota was performed to determine if there were significant relationships between organism size and MP abundance. There was no correlation between the length of individual organisms and abundance of MPs (*R*^*2*^ = 0.00, *F*(1,255) = 0.49, p = 0.4857), but the weight of individual organisms was significantly correlated to the number of MP per individual (*R*^*2*^ = 0.07; *F*(1, 283) = 21.68, p < .0001) (**Figure 2**).

**Figure 1.**
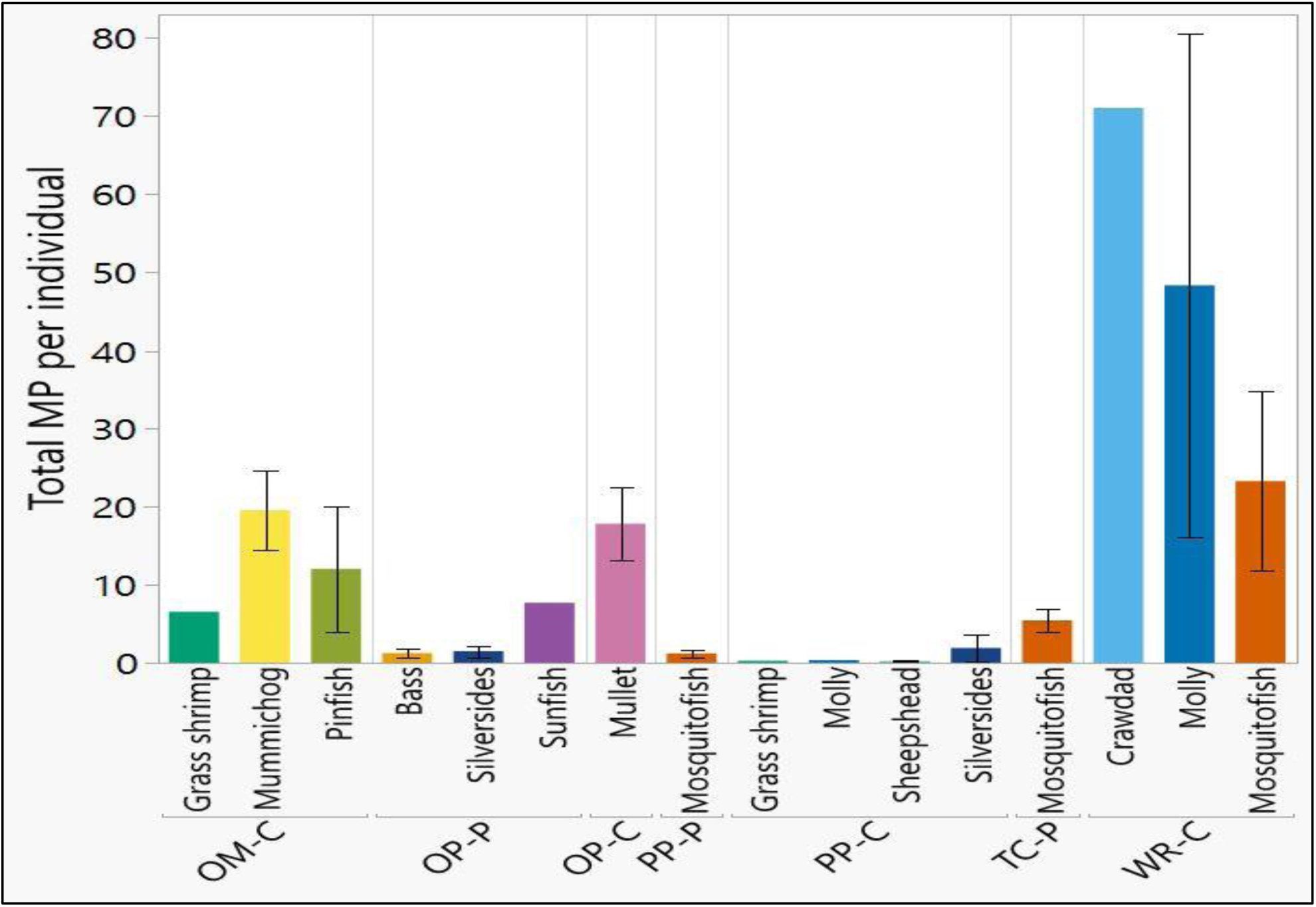
Average total microplastics per individual by site and species. Total microplastics includes fibers, fragments, and suspected tire particles. Whiskers indicate standard error. The absence of whiskers indicates samples where standard error could not be calculated due to small sample size. OM-C is Oak Marsh creek, OP-P is Oyster Point pond, OP-C is Oyster Point creek, PP-P is Patriots Point pond, PP-C is Patriots Point creek, TC-P is Tides Condos pond, and WR-C is Whipple Road creek.

**Figure 2.**
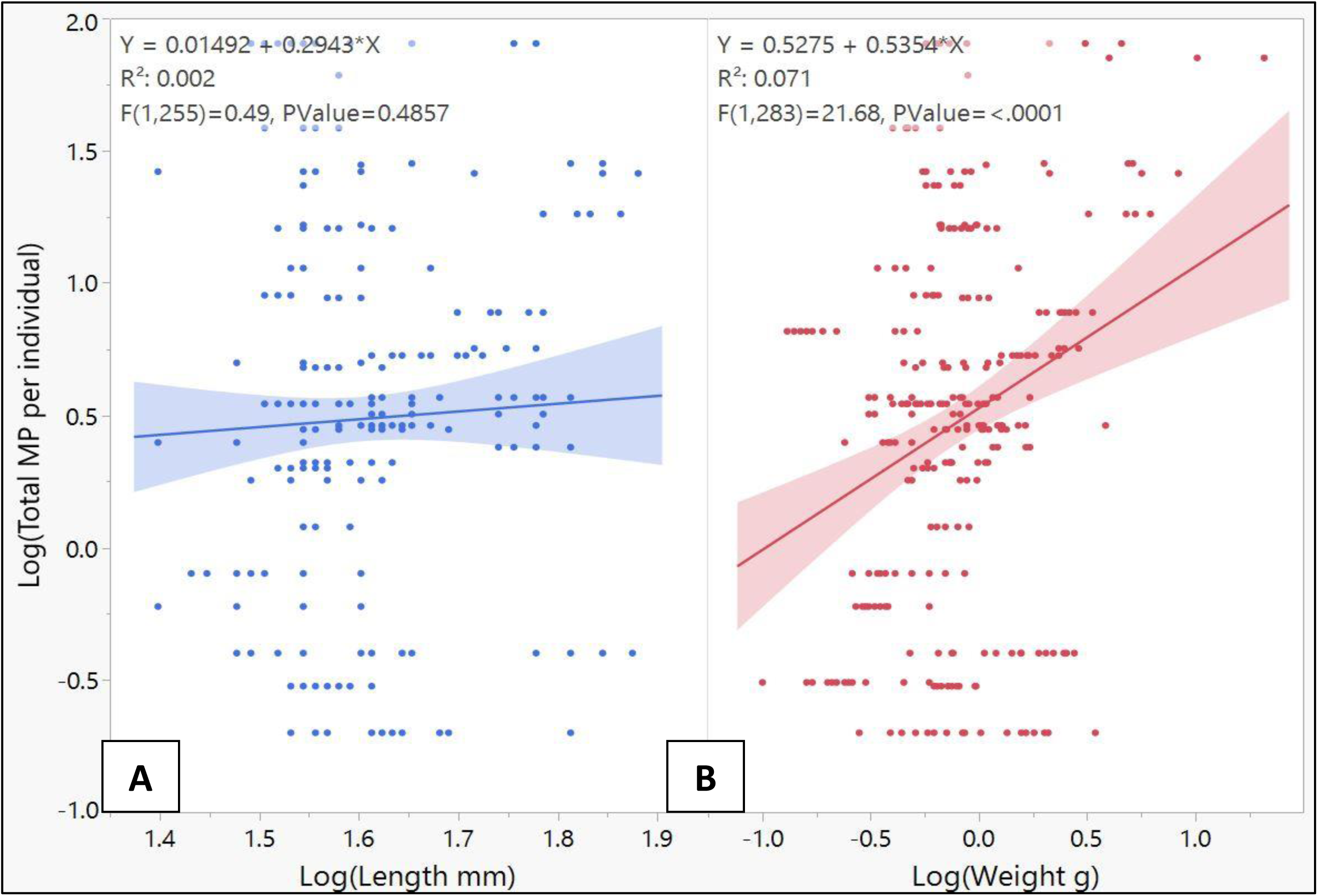
Length in mm (A) and weight in g (B) of individual organisms and average microplastics (including fibers, suspected tire particles, and fragments) per individual. The shaded region represents the 95% confidence interval.

Given the significant relationship observed between the number of MP per individual and weight, MP abundance was also analyzed in relation to organism weight for comparisons between sites and between species (**Figure 3**).

**Figure 3.**
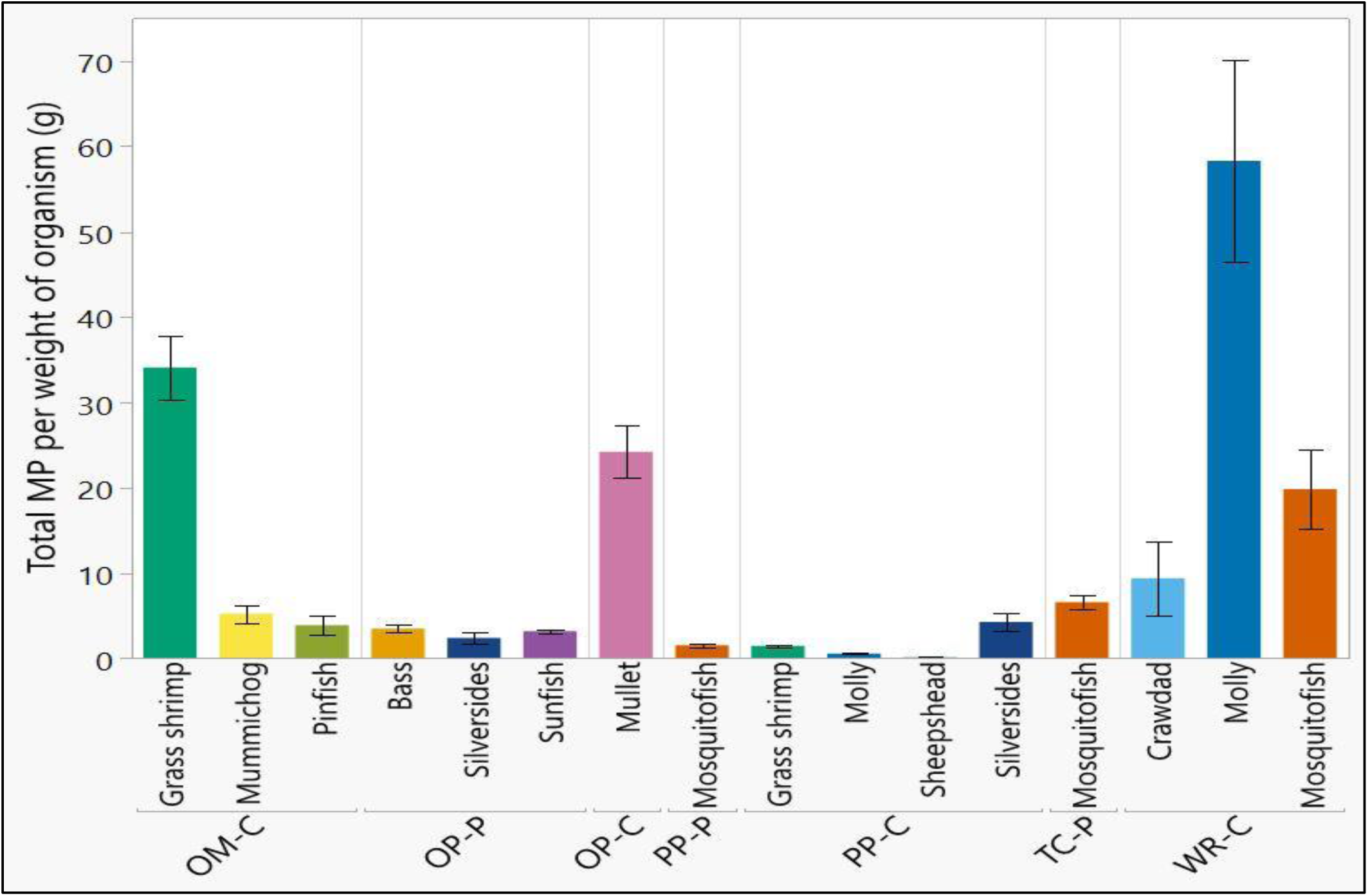
Average total microplastics by weight (gram) of organism by site and species. Total microplastics includes fibers, fragments, and suspected tire particles. Whiskers indicate standard error. The absence of whiskers indicates samples where standard error could not be calculated due to small sample size. OM-C is Oak Marsh creek, OP-P is Oyster Point pond, OP-C is Oyster Point creek, PP-P is Patriots Point pond, PP-C is Patriots Point creek, TC-P is Tides Condos pond, and WR-C is Whipple Road creek.

Microplastics were classified by size fraction according to sieve mesh size and were either 53µm – 500µm or 500µm – 5mm in size. Of the 2,713 MP counted, 74.8% were between 53µm - 500µm and 25.1% were 500 µm – 5mm in size. There were significantly more MP in the size range from 53µm – 500µm (*t*(58) = -2.32, p = 0.0119). Microplastics were also categorized into three major types: fibers, suspected tire particles (TP) or fragments. (**Figure 4**).

**Figure 4.**
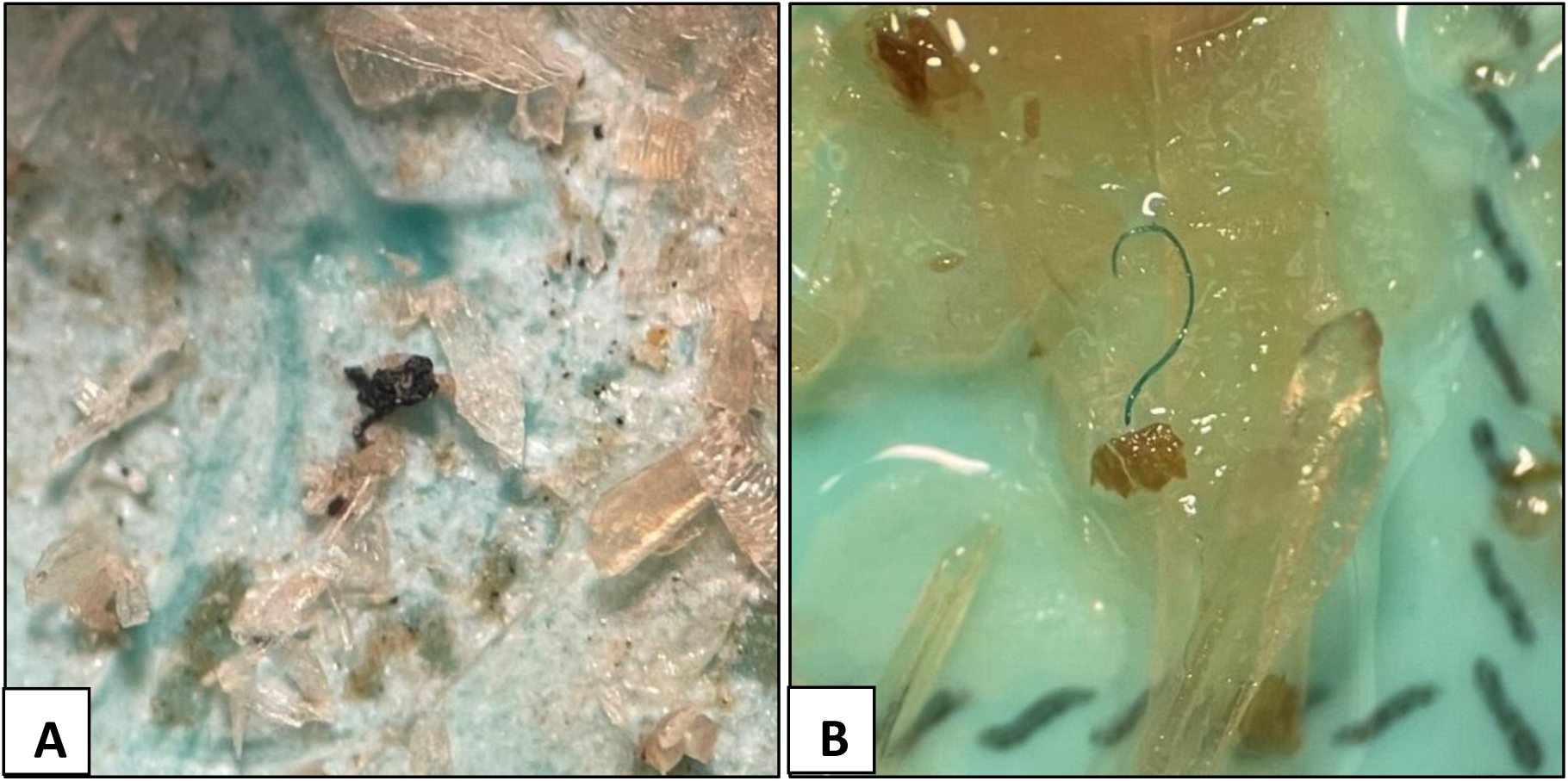
Microplastic types in biota from stormwater pond and adjacent tidal creek sites. (A) A suspected tire particle and (B) a blue fiber.

The distribution of MP types encountered was 8.0% fibers, 89.9% suspected tire particles, and 2.1% fragments across all MPs counted. The focus of this study was understanding the occurrence of suspected tire wear particles in organisms from stormwater ponds and their adjacent tidal creeks; therefore, the microplastics were divided into two groups for further analysis, suspected tire particles (TP) and fibers + fragments. The distribution of MP types across sites indicated that suspected tire particles consistently made up the majority of total MP (**Figure 5**).

**Figure 5.**
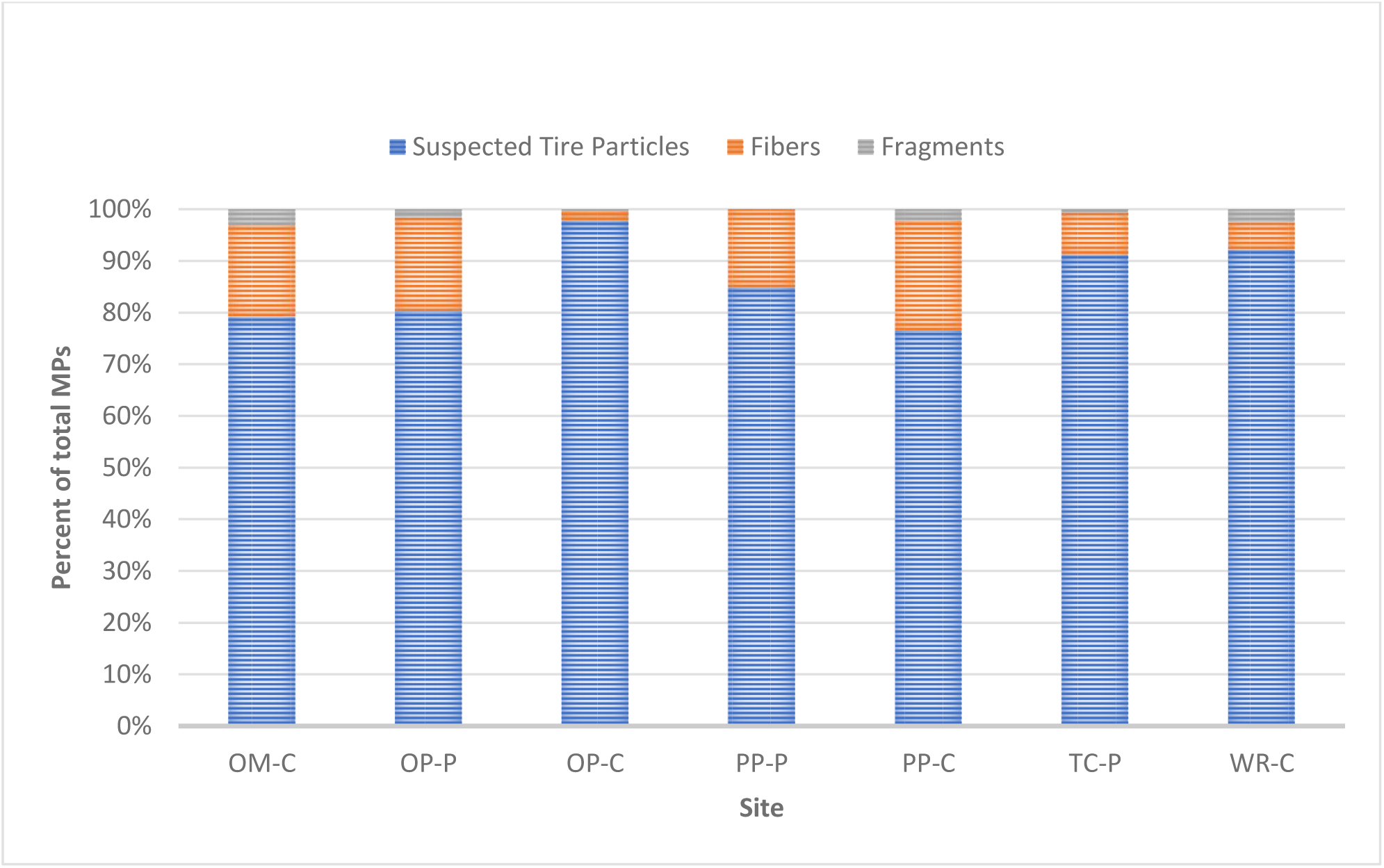
Distribution of microplastic types in organisms collected from stormwater ponds and adjacent tidal creek sites. OM-C is Oak Marsh creek, OP-P is Oyster Point pond, OP-C is Oyster Point creek, PP-P is Patriots Point pond, PP-C is Patriots Point creek, TC-P is Tides Condos pond, and WR-C is Whipple Road creek.

### Site differences

Total microplastics (fibers, suspected tire particles, fragments) per individual were compared across all sites sampled (n = 7 sites), regardless of species or size fraction (**Figure 6, Table 1**). There was a significant difference in total MP per individual among sites (*F*(6, 37) = 11.99, p = <.0001). The number of MP per individual was significantly greater at the Whipple Road creek site (WR-C, 35.5 ± 11.2 MP per individual) site compared to Patriots Point pond (PP-P, 1.3 ± 0.5 MP per individual, and Patriots Point creek (PP-C, 0.9 ± 0.6 MP per individual), and Oyster Point pond (OP-P, 2.2 ± 0.9 MP per individual) (Tukey’s HSD, p = 0.0003, p = < .0001, and p = 0.0003 respectively).

**Table 1.** Microplastic counts and types in each species across all sites sampled. Total microplastics includes fibers, fragments, and suspected tire particles. n = number of individuals collected at each site.

| Site | n | Total microplastics per individual (mean ± SEM) | Tire particles per individual (mean ± SEM) |
| --- | --- | --- | --- |
| Whipple Road Creek (WR-C) |  |  |  |
| Crawdad | 3 | 71 | 57.7 |
| Molly | 19 | 48.3 ± 32.2 | 48.2 ± 32.2 |
| Mosquitofish | 36 | 23.3 ± 11.5 | 17.6 ± 8.4 |
| *Overall for site* | 58 | 35.5 ± 11.2 | 30.3 ± 10.1 |
| Tides Condos Pond (TC-P) |  |  |  |
| Mosquitofish | 30 | 5.5 ± 1.5 | 5.0 ± 1.3 |
| *Overall for site* | 30 | 5.5 ± 1.5 | 5.0 ± 1.3 |
| Patriots Point Pond (PP-P) |  |  |  |
| Mosquitofish | 30 | 1.3 ± 0.5 | 1.1 ± 0.4 |
| *Overall for site* | 30 | 1.3 ± 0.5 | 1.1 ± 0.4 |
| Patriots Point Creek (PP-C) |  |  |  |
| Grass shrimp | 13 | 0.3 | 0.2 |
| Molly | 5 | 0.4 | 0 |
| Sheepshead | 10 | 0.3 ± 0.1 | 0 |
| Silversides | 15 | 2.0 ± 1.8 | 1.7 ± 1.6 |
| *Overall for site* | 43 | 0.9 ± 0.6 | 0.6 ± 0.5 |
| Oyster Point Pond (OP-P) |  |  |  |
| Bass | 30 | 1.3 ± 0.6 | 1.0 ± 0.4 |
| Silversides | 20 | 1.6 ± 0.7 | 1.1 ± 0.7 |
| Sunfish | 8 | 7.8 | 6.8 |
| *Overall for site* | 58 | 2.2 ± 0.9 | 1.8 ± 0.8 |
| Oyster Point Creek (OP-C) |  |  |  |
| Mullet | 21 | 17.9 ± 4.8 | 17.5 ± 4.6 |
| *Overall for site* | 21 | 17.9 ± 4.8 | 17.5 ± 4.6 |
| Oak Marsh Creek (OM-C) |  |  |  |
| Grass shrimp | 12 | 6.6 | 4.58 |
| Mummichog | 13 | 19.6 ± 5.1 | 16.2 ± 5.4 |
| Pinfish | 20 | 12.1 ± 8.0 | 9.6 ± 6.7 |
| *Overall for site* | 45 | 15.1 ± 4.0 | 12.3 ± 3.7 |

**Figure 6.**
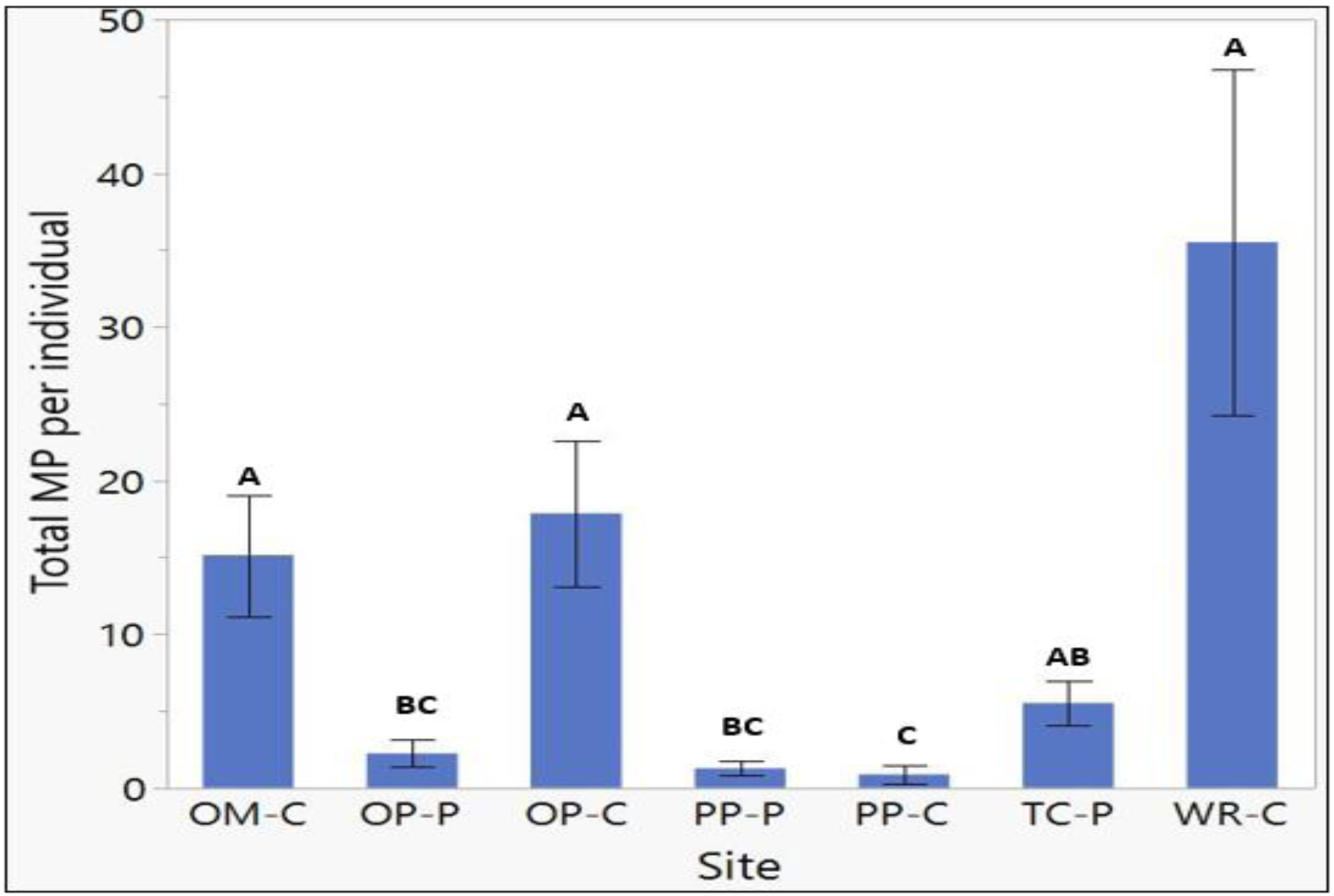
Average total microplastics (fibers, fragments, suspected tire particles) per individual collected from stormwater ponds and adjacent tidal creek sites. Whiskers indicate standard error. Different letters indicate significant difference between sites (p < 0.05). OM-C is Oak Marsh creek, OP-P is Oyster Point pond, OP-C is Oyster Point creek, PP-P is Patriots Point pond, PP-C is Patriots Point creek, TC-P is Tides Condos pond, and WR-C is Whipple Road creek.

There were also significantly more total MP per individual in the Oak Marsh creek site (OM-C, 15.1 ± 4.0 MP per individual) compared to Patriots Point creek, Patriots Point pond, and Oyster Point pond (Tukey’s HSD, p = < .0001, p = 0.0033, and p = 0.0055, respectively).

The Oyster Point creek site (OP-C, 17.9 ± 4.8 MP per individual) had significantly more MP per individual when compared to the reference sites (Patriots Point creek and Patriots Point pond) and when compared to the pond site for the same location, Oyster Point pond (Tukey’s HSD, p = 0.0003, p = 0.0058, and p = 0.0114, respectively). Lastly, the Tides Condos pond site (TC-P, 5.5 ± 1.5 MP per individual) had significantly more MP per individual compared to the reference creek site, Patriots Point creek (Tukey’s HSD, p = 0.0172).

Data were also analyzed as total MP per body weight (g) of organism. There was a significant difference in MP abundance per g among sites (*F*(6,278) = 47.52, p = < 0.0001). When normalized to weight of the organism, differences in MP abundance between sites were more apparent with the reference sites at Patriots Point having significantly fewer MP compared to the other sampling sites (**Figure 7)**.

**Figure 7.**
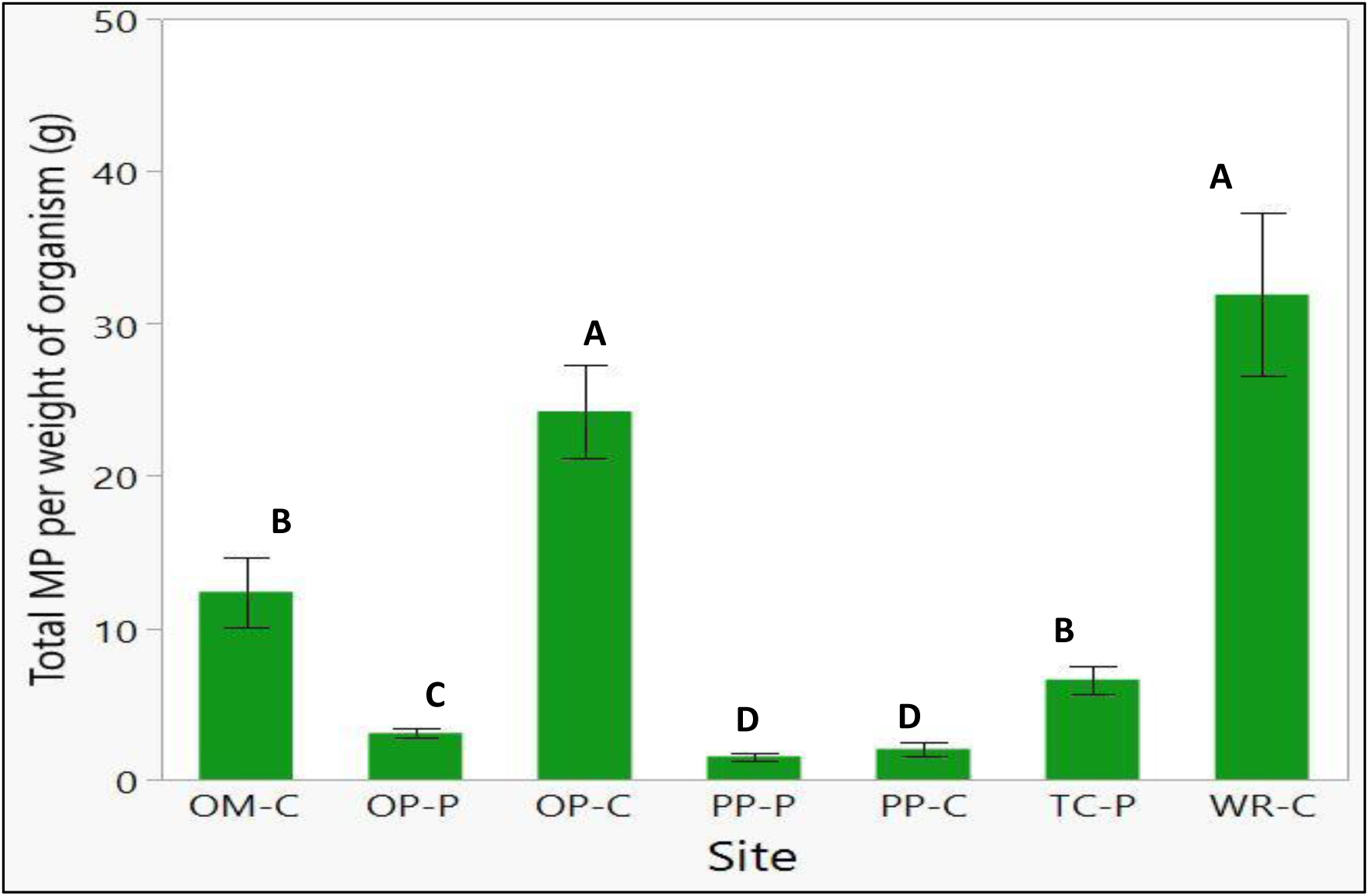
Average total microplastics (fibers, fragments, suspected tire particles) by weight (g) in organisms collected from stormwater ponds and adjacent tidal creek sites. Whiskers indicate standard error. Different letters indicate significant difference between sites (p < 0.05). OM-C is Oak Marsh creek, OP-P is Oyster Point pond, OP-C is Oyster Point creek, PP-P is Patriots Point pond, PP-C is Patriots Point creek, TC-P is Tides Condos pond, and WR-C is Whipple Road creek.

There were only two locations for which samples from both the pond and adjacent tidal creek were collected: the reference site, Patriots Point, and Oyster Point. At Patriots Point, the total MPs per individual were not significantly different between the tidal creek (PP-C, 0.9 ± 0.6 MP per individual) and the pond (PP-P, 1.3 ± 0.5 MP per individual) (*t*(8.4) = 0.907, p = 0.3892) (**Figure 8A**). At Oyster Point, there were significantly more total MPs per individual in the creek (OP-C, 17.9 ± 4.8 MP per individual) compared to the pond (OP-P, 2.2 ± 0.9 MP per individual) (*t*(9.2) = -4.325, p = 0.0009) (**Figure 8B**). This was unexpected assuming that the ponds contain more MP in general and thus would result in more MP in organisms collected from the ponds compared to the creeks.

**Figure 8.**
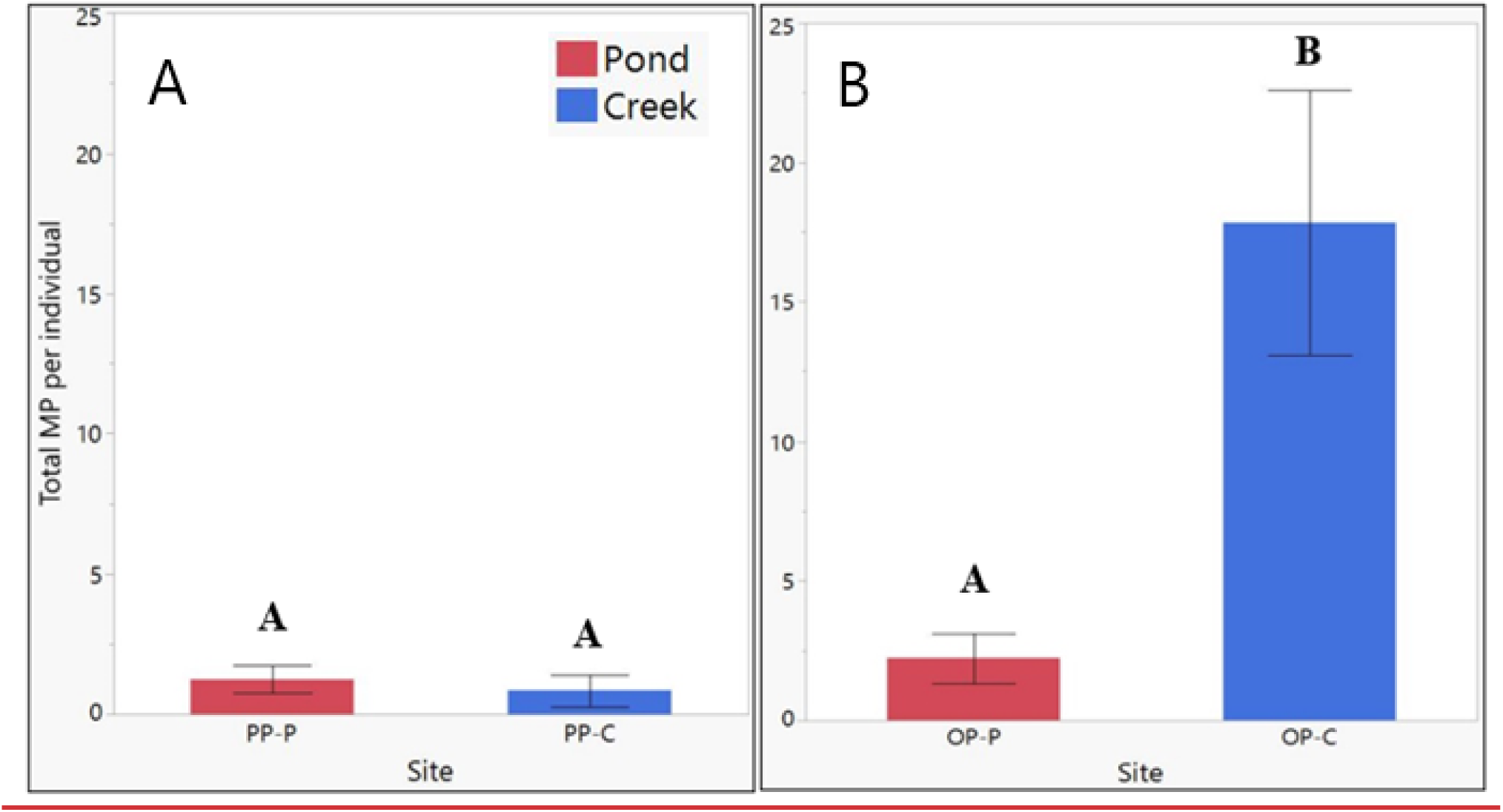
**A,B**Average total microplastics (fibers, fragments, suspected tire particles) per individual collected from stormwater ponds (P) and adjacent tidal creek (C) sites for Patriots Point (PP) and Oyster Point (OP) locations. Whiskers indicate standard error. Different letters indicate significant difference between sites (p < 0.05).

### Species differences

To assess species differences in microplastic abundance in biota, microplastics per individual were compared across all organisms collected, regardless of site or size fraction (**Figure 9, Table 1**). There was a significant difference in total MP per individual among species (*F*(10, 274) = 22.55, p = < 0.0001). Tukey’s HSD post-hoc analysis for comparisons of each pair showed there were significantly more MP per individual in crawdads (71 MP per individual) and mollies (48.3 ± 32.2 per individual) compared to pinfish, mosquitofish, grass shrimp, silversides, bass, and sheepshead. There were significantly more MP per individual in mummichogs (19.6 ± 5.1 MP per individual) and mullet (17.9 ± 4.8 MP per individual) compared to mosquitofish, grass shrimp, silversides, bass, and sheepshead. Sunfish (7.8 MP per individual), pinfish (12.1 ± 8.0 MP per individual), and mosquitofish (10.0 ± 3.3 MP per individual) had significantly more MP per individual than silversides, bass, and sheepshead. Sheepshead minnow total MP per individual (0.3 ± 0.1 MP per individual) were significantly lower than all species except silversides and bass.

**Figure 9.**
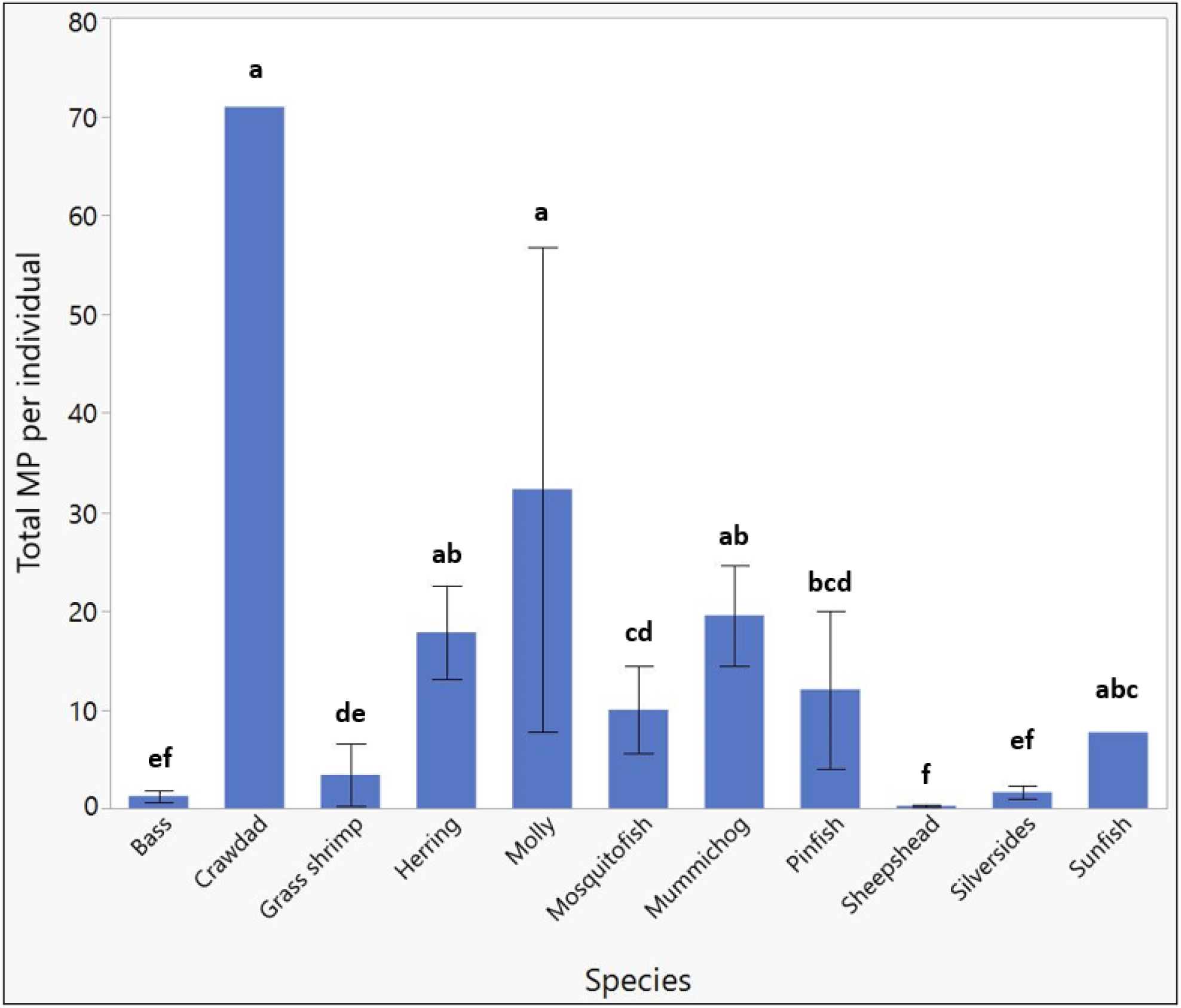
Average total microplastics per individual collected from stormwater ponds and adjacent tidal creek sites for each species collected. Whiskers indicate standard error. The absence of whiskers indicates samples where standard error could not be calculated due to small sample size. Different letters indicate significant difference between species (p < 0.05).

Data were also analyzed as total MP per weight (g) of organism. There was a significant difference in MP abundance per g among species (*F*(10,274) = 15.55, p = < 0.0001) (**Figure 10**). When normalized to weight of the organism, differences in MP abundance between species seem to suggest other variables such as feeding habitat may be responsible for the observed differences. When making comparisons among species, it is important to acknowledge that not all species were collected at each site. For example, sheepshead were only collected from the reference site creek (PP-C) and therefore have the lowest MP abundance recorded.

**Figure 10.**
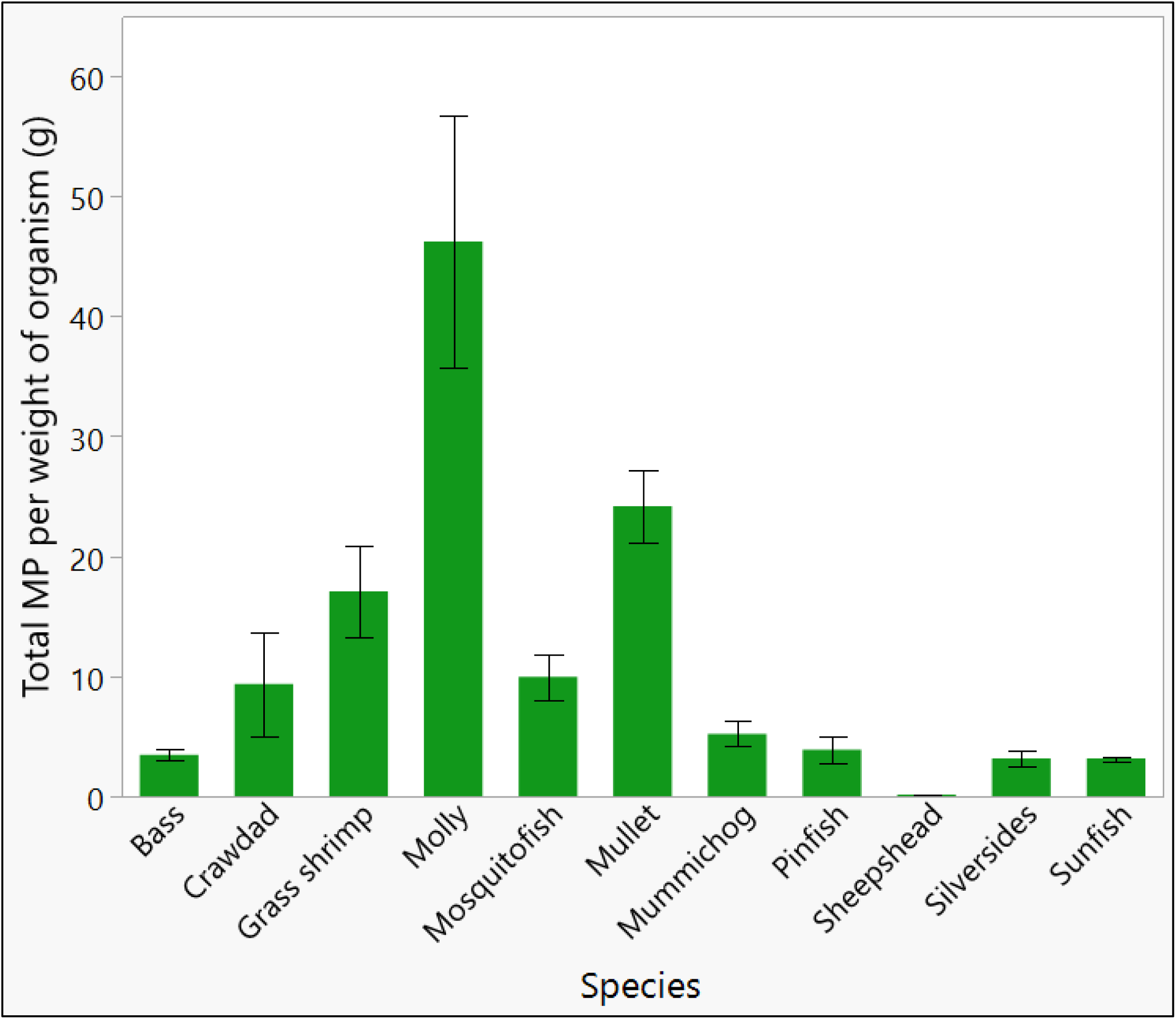
Average total microplastics by weight (g) of organisms collected from stormwater ponds and adjacent tidal creek sites for each species collected. Whiskers indicate standard error. Different letters indicate significant difference between species (p < 0.05).

## Discussion

### Microplastic counts – size fraction recovered and distribution of MP types recovered

Biota collected from stormwater pond and adjacent tidal creeks in Mount Pleasant, SC, USA were found to contain an average of 9.5 ± 6.5 MP per individual, which included fibers, suspected tire particles, and fragments. The organisms analyzed included nine species of fish, one species of grass shrimp, and one species of crawdad (**Table S2**). Other studies on microplastic abundance in biota mostly focused on marine species or freshwater taxa. The range of MPs reported in biota can vary depending on several factors including species, geographic location, habitat, anthropogenic influence and nearby land use, or seasonal weather patterns. The average number of MP per fish from the present study were within the range of MP concentrations observed in other studies; for example, 9.9 ± 13.4 particles per fish were found in organisms collected upstream and downstream from a sewage treatment plant in Korea (Park et al., 2020b). Additionally, other estimates of MP in various species and geographic locations include 0.19 to 1.63 MP per individual for *Lepomis* spp. from the Brazos River Basin, USA (Peters &Bratton, 2016), 0 to 18 MP per individual in *Carassius auratus* collected from a freshwater lake in China (Yuan et al., 2019), an average of 1.71 ± 2.27 MP per individual in the terrestrial crab *Cardisoma carnifex* from Vanuatu (Bakir et al., 2020), and an average of 1.8 ± 1.7 MP per individual in coastal and offshore fish from the Northeast Atlantic (Murphy et al., 2017). The lack of standard reporting for microplastic identification and sample processing of microplastics in biota made it difficult to compare studies as both microplastic identification methods and sample processing methods (i.e., gut only vs. whole organism) can vary. Nevertheless, the reported ranges for MPs in biota from freshwater environments were typically greater than those observed in marine environments, and a general reduction in the abundance of MPs from land (i.e., freshwater) to nearshore to offshore has been observed suggesting that anthropogenic influence greatly impacts MP abundance in different environmental matrices (Bakir et al. 2020; Graca et al., 2017).

Parker et al. (2020) assessed microplastics in estuarine fishes in the same geographical area as this study (Charleston Harbor, SC, USA), and found an average of 26.9 ± 4.7 MP per fish. Fish processing methods (digestion of whole organism in KOH) and the size range of MPs analyzed (63 µm to 5mm) were similar to the present study, allowing for good comparability between studies. They observed a much higher average number of MP per individual than was observed in the present study, which may be attributed to the larger size of the fish collected and analyzed. The authors suggested, along with others, that MP abundance increases as fish size increases (Hossain et al., 2019; Hurt et al., 2020; Parker et al., 2020; Peters &Bratton, 2016). When normalized to the weight of fish, Parker et al. (2020) observed an average of 5.8 ± 1.6 MP per g fish, which is comparable to the present study, where we observed an average of 3.87 MP per g body weight. Using length of fish, Parker et al. (2020) reported an average length of 104 ± 6 (mm), therefore, an average of 0.25 MP per mm when normalized to MP per length of fish. The present study found an average of 0.31 MP per mm fish (crawdads and grass shrimp not included).

Data available from studies that assess MP abundance in the water column, sediment, and biota suggested that in general, MPs accumulate the most in sediments followed by the water column, then biota, although local differences can occur depending on environmental mixing and flow and species examined (Cera et al., 2020; Kazour et al., 2019; Yuan et al., 2019; Zhang et al., 2020). For stormwater ponds specifically, only one study to date has collected samples for MP analysis from the water column, sediment, and biota and found that MPs accumulated to the highest concentrations in sediments, followed by vertebrates analyzed from the pond, with the water column having the least number of MPs (Olesen et al., 2019). Although different water bodies were examined compared to the stormwater ponds examined in the present study, MP abundances in the Charleston Harbor estuary indicated that MPs in the water ranged from 3 to 36 MP/m^2^ in water and 0 to 4,375 MP per kg wet weight in sediments (Leads &Weinstein, 2019).

Additionally, for biota, Payton et al. (2020) observed 1.4% of zooplankton collected from the Cooper River front (adjacent to Mt. Pleasant, SC, USA) to contain microplastics.

However, determining the MP abundance in stormwater ponds in the Charleston area is currently in its infancy. Kell (2020) reported greater MPs in stormwater pond water and sediment compared to MPs in adjacent discharge creek water and sediment for stormwater ponds also located on Mt. Pleasant, SC USA. The data presented here reflect a different trend, namely that there were more MPs in biota from adjacent tidal creeks compared to MPs in biota from stormwater ponds. However, this observation is somewhat inconclusive as there was only one sampling location, besides the reference site, where organisms were collected from both the stormwater pond and tidal creek. For stormwater ponds globally, observations show a greater amounts of MPs in stormwater pond sediment and water compared to discharge point samples (**Table 2)**. These studies show that regardless of size fraction analyzed, stormwater pond sediments seem to act as significant sinks for microplastics. Additional data on the abundance of MP in the water, sediment, and adjacent tidal creek water and sediment from the stormwater ponds in the present study are being collected and will assist in forming conclusions regarding the availability of MP to stormwater pond biota.

**Table 2.** Microplastic abundances in studies from Charleston Harbor, SC area and stormwater pond studies worldwide. Microplastics are reported as averages or min and max averages unless otherwise noted.

| Sample type | Location | Size range | Microplastics (Average) | Study |
| --- | --- | --- | --- | --- |
| **Charleston Harbor, SC, USA studies** |  |  |  |  |
| Surface water | Charleston, SC USA | > 63 µm | 3 – 36 MP L<sup>-1</sup> (min – max) | Leads & Weinstein, 2019 |
| Sediment | Charleston, SC USA | > 63 µm | 0 – 4,375 MP kg<sup>-1</sup> ww<sup>-1</sup> (min – max) | Leads & Weinstein, 2019 |
| Surface water | Charleston, SC USA | > 63 µm | 3 – 11 MP L<sup>-1</sup> (min – max) | Gray et al., 2018 |
| Sediment | Charleston, SC USA | > 63 µm | 42.2 – 11,195.7 MP m<sup>-2</sup> (min – max) | Gray et al., 2018 |
| Surface water | Charleston, SC USA | 43 - 104µm | 0.6 – 1.0 MP L<sup>-1</sup> | Payton et al., 2020 |
| Biota (zooplankton) | Charleston, SC USA | 43 - 104µm | avg. 1.4% of individuals | Payton et al., 2020 |
| Biota (fish) | Charleston, SC USA | 43µm – 11.3mm | 27 MP individual<sup>-1</sup> | Parker et al., 2020 |
| **Stormwater pond studies** |  |  |  |  |
| Pond water | Charleston, SC USA | 63 - 5000µm | 1.6 – 132 MP L<sup>-1</sup> (min – max) | Kell, 2020 |
| Pond sediment | Charleston, SC USA | 63 - 5000µm | 1,379 - 10,557 MP kg<sup>-1</sup> dw<sup>-1</sup> (min – max) | Kell, 2020 |
| Tidal creek sediment adjacent to stormwater pond | Charleston, SC USA | 63 - 5000µm | 100 – 10,000 MP kg<sup>-1</sup> dw<sup>-1</sup> (min – max) | Kell, 2020 |
| Pond water | Denmark | 10 - 500µm | 270 MP L<sup>-1</sup> | Olesen et al., 2019 |
| Pond sediment | Denmark | 10 - 500µm | 9.5 x 10<sup>5</sup> MP kg<sup>-1</sup> dw<sup>-1</sup> | Olesen et al., 2019 |
| Pond biota | Denmark | 10 - 500µm | 65 MP individual<sup>-1</sup> | Olesen et al., 2019 |
| Pond water | Denmark | 10 - 2000µm | 490 – 22,894 MP m<sup>-3</sup> (min – max) | Liu et al., 2019 |
| Pond sediment | Denmark | 10 - 2000µm | 1,115 – 128,732 MP kg<sup>-1</sup> dw<sup>-1</sup> (min – max) | Liu et al., 2019 |
| Pond water | Sweden | 20 - 300µm | 977 – 4,267 MP L<sup>-1</sup> (min – max) | Jönsson, 2016 |
| Pond water | Australia | 25 - 500µm | 0.9 – 4.0 MP L<sup>-1</sup> (min – max) | Ziajahromi et al., 2020 |
| Pond sediment | Australia | 25 - 500µm | 320 – 595 MP kg<sup>-1</sup> dw<sup>-1</sup> (min – max) | Ziajahromi et al., 2020 |

In the present study, total MPs were classified by two size fractions: 53µm – 500µm and >500µm. There were significantly more MP collected in the smaller size range, which is corroborated by other studies that also found a greater abundance of the smaller sized MPs in surface waters, sediments, and biota (Bakir et al., 2020; Olesen et al., 2019; Park et al., 2020a; Park et al., 2020b; Ziajahromi et al., 2020). Smaller particles are potentially more prevalent in biota collected from stormwater ponds because i) they closely align with the size range of natural food items for the biota captured (Parker et al., 2020) and ii) larger particles may effectively be trapped or settled in sediments and less likely to be consumed (Besseling et al., 2017).

The majority (>80%) of total MPs counted in individuals across all sites were suspected tire particles, highlighting the importance of stormwater ponds and their discharge points into tidal creeks as pathways and potential hot spots for MR in the environment. The majority of tire wear particles collected from the environment may range between 25 and 50 µm in size (Kreider et al.. 2010) which was below the threshold for the size sampled in this study (53 µm). Although the majority of total MP counted in the present study were suspected tire particles, difficulty with confirming polymer type for suspected tire particles may have led to inaccurate counts. Nevertheless, other studies have documented suspected tire particles (25 - >500µm) in sediment samples, comprising up to 38% of total MPs, from both inlets and outlets of stormwater ponds, indicating they are available in stormwater ponds and at discharge points (Ziajahromi et al., 2020).

Additionally, Kell (2020) reported high abundances of TP in stormwater pond sediment and sediment of pond discharge points, up to 80% of total MP in pond sediments and up to 60% of total MP in discharge point sediments and more specifically, the majority were between 63 - 500µm in size. These data suggest that TP are accumulating in stormwater ponds from stormwater runoff and adjacent tidal creeks where ponds discharge and are available to biota in stormwater ponds and their adjacent tidal creeks.

### Site and species differences

The average number of MP per individual was significantly different among the sampling sites. The reference tidal creek, Patriots Point creek, contained significantly fewer MP per individual (0.9 ± 0.6 MP per individual) compared to all other sites. The reference pond site, Patriots Point pond, also contained significantly fewer MP per individual (1.3 ± 0.5 MP per individual) than Whipple Road, Oyster Point, and Oak Marsh tidal creek sites. Interestingly, there was not a significant difference between the average number of MP per individual in Patriots Point pond compared to the other pond sites sampled, Oyster Point and Tides Condos ponds. Of those two pond sites, Oyster Point pond had significantly fewer MP per individual compared to its adjacent tidal creek (OP-C) as well as Oak Marsh creek and Whipple Road creek whereas Tides Condos pond had significantly more MP per individual but only when compared to the reference tidal creek, PP-C.

Based on the high approximate daily traffic from traffic monitoring locations in proximity to the ponds and tidal creek sites (**Table S1**), it was hypothesized that Tides Condos pond (662,700 vehicles day^-1^) would have the greatest number of MP per individual, especially for suspected tire particles. However, this hypothesis was not supported in the present study, and instead, Whipple Road creek was found to contain the greatest amount of MP per individual collected. Whipple Road sites also experienced contributions from roadway pollution and although the area has lower daily traffic counts (11,500 vehicles day^-1^) compared to other sites sampled, this site was directly adjacent to a major throughfare street, Whipple Road. Liu et al. (2019) evaluated MP abundance in the water phase from stormwater ponds that drained different land use areas including residential, industrial, commercial, and highway areas and found a greater abundance of MPs in stormwater ponds that drained industrial and commercial land areas compared to ponds serving highway and residential areas; however, their analysis did not include car tire rubber, which would be a major contributor to stormwater MP in ponds draining highway runoff.

The lower MP collected per individual from Tides Condos pond may be due to the fact that at this pond only one species (mosquitofish) was collected and analyzed. This could result in an inaccurate characterization of the abundance of MP in overall pond biota because the sample was not representative of the entire pond population. Data for MP abundance in biota from Whipple Road pond could not be established for the present study, but based on observations from Kell (2020) and others regarding MP abundance in sediments of stormwater ponds and discharge points, we expect that the high abundance of MP in Whipple Road creek is due to a combination of site-specific attributes that allow for relatively large discharge of MP into the adjacent tidal creek of this pond. The MTD associated with the Whipple Road site serviced a large watershed catchment that included several neighborhoods, a large church parking lot, and the two-lane throughfare street, Whipple Road. Additionally, the MTD at this site was flawed in that it experienced back flow of water into the MTD during large storm events or very high tides. The MTD’s outflow pipe was flush with the creek, allowing for a more direct connection between the pond and adjacent tidal creek.

The other two tidal creek sites sampled with relatively high abundances of MP were Oyster Point and Oak Marsh creek. The Oyster Point locations were representative of new (< 10 years old) residential areas. Surprisingly, Oyster Point pond contained less MP per individual compared to Oyster Point creek. Species diversity and heterogeneity of samples may have influenced MP abundances between these two sites where the sample from the pond had bass, silversides, and sunfish and the sample from the creek only had mullet. Differences in foraging behavior or preferred habitat between species collected may partially explain the differences in average MPs collected from the two sites.

Juvenile bass, silversides, and sunfish are omnivorous feeders throughout the water column whereas mullet are primarily detritivores that feed closer to the bottom (Antonucci et al., 2014; Bester, 2017; Carlander, 1977; Miranda &Pugh, 1997). McNeish et al. (2018) argued that species traits can help explain microplastic abundance and these are species dependent. They found that MP abundance was positively related to fish trophic level, where zoobenthivores had greater MP abundance compared to omnivores.

Oak Marsh creek had the third highest average MP per individual of all sites sampled. Oak Marsh pond receives stormwater from residential and highway areas, with daily traffic based on traffic monitoring locations in proximity to the Oak Marsh sites estimated at 49,000 vehicles daily. Additionally, influx likely included airborne particulate contamination from a major highway, Intestate-526, adjacent to the Oak Marsh sites. Therefore, we hypothesized there would be a high amount of TP at the Oak Marsh sites, which was indeed observed (79% of all MP at OM-C). Data on the abundance of MP in biota from Oak Marsh pond was unavailable for comparison because of site specific sampling problems. Species collected at Oak Marsh creek included grass shrimp, mummichog, and pinfish and represented benthic omnivores, epibenthic omnivores, and carnivorous species, respectively (Abraham, 1985; Feinstein, 1975; Odum &Heald, 1972). The observed positive correlation between biota weight and MP abundance may also explain the high abundance of MP in biota from Oak Marsh creek as mummichogs were the second-largest species collected (behind crawdads) at 4.97 ± 2.15 g body weight, and pinfish were also one of the larger species collected at 1.57 ± 0.14 g. Oak Marsh creek specimens were collected during a king tide with exceptionally high-water levels, creating backflow from the creek into the pond. It is possible that the tidal influence caused resuspension of MPs from the pond and creek sediments which could have contributed to the overall greater MP per individual observed at this site.

As for species differences in MP abundance, there were significant differences in total MP per individual among species collected. Crawdads had the most MP with an average of 71 MP per individual. Crawdads represent one of the truly benthic organisms collected in this study. A greater number of MPs were expected in bottom-dwelling organisms that interact directly with sediment that contains settled particles. In comparison, grass shrimp which are epibenthic feeders but also feed in floating vegetation, contained significantly less MP per individual (3.4 ± 3.1) than crawdads, but this difference may be attributed to the smaller size of grass shrimp and differences in size of natural food items typically ingested by each species.

As for fish species, there was a wide variability in MP abundance. As previously stated, size, trophic position, feeding strategy, or habitat may influence the distribution of MPs among species. Park et al. (2020a) found more MP per individual in bottom dwelling omnivorous fish (i.e. carp) compared to epibenthic and pelagic omnivorous and carnivorous fish (minnow and bass, respectively). While differences were observed between species in the present study, there were no clear relationships between MP per individual and species, likely due to a combination of site-specific availability of MP, organism size, and feeding habitat influencing the total MP observed.

## 4.6 Conclusions

Stormwater ponds can function as effective stormwater runoff best management practices and trap or remove both physical and chemical pollutants before further discharge into receiving natural waterbodies. Stormwater ponds can therefore potentially accumulate high levels of pollutants, including microplastics and microrubber from road runoff. It was hypothesized that biota in stormwater ponds in proximity to roadways or those that receive large road runoff would contain relatively high amounts of MP per individual, with a high abundances of TP. Indeed, the data indicated that the majority of MP recovered from biota across all sites were suspected TP.

There were significant differences in MP per individual observed between sites and between species. It seemed that a combination of factors such as availability of MP, organism size, and feeding habitat or behavior influenced the total MP observed. A significant positive correlation between MP per individual and organism weight was observed, with larger individuals typically containing more MP per individual. It appeared that species-specific feeding habits influence the total MP observed, as was found for some benthic and sediment-dwelling organisms who are exposed to sediments that contain more available MP compared to pelagic dwelling organisms. However, examining the detailed niche of each species in relation to MP abundance was not within the scope of the present study.

Comparisons between pond and adjacent tidal creek sites were somewhat inconclusive. There were only two locations in which both the pond and the tidal creek could be sampled. Of these two, one was the reference site (Patriots Point) which had no significant difference between total MP per individual between the pond and creek.

Additionally, MP abundance from Patriots Point creek was significantly lower compared to all other sites (except Patriots Point pond), making it a good reference site for MP abundance in biota. For the other sites, it was hypothesized that there would be more MP, specifically more TP, in biota from pond sites compared to creek sites. The results are inconclusive due to the small sample size and inability to collect organisms from all ponds and all tidal creeks. Completing the dataset by capturing biota from all ponds and all tidal creeks and analyzing sediment and water column samples for MPs will assist with making more accurate comparisons between the two and determining the efficiencies of stormwater ponds in retaining MP. Future work should attempt to collect and analyze the same species from both pond and tidal creeks or species with similar feeding mechanisms and habitat preferences for better comparability.

A major limitation of this study was the exact identification of MP, specifically suspected tire wear particles. Additional analysis of a subset of MP and suspected TP using advanced spectroscopy such as micro-attenuated total reflectance (ATR) Fourier transform infrared (FTIR) spectroscopy for MPs (< 500µm) or scanning electron microscopy (SEM) for TPs would be beneficial to confirm suspected tire wear particle identity. Tire particles can be analyzed by ATR-FTIR and µATR-FTIR methods but the presence of filler materials such as carbon black confound analytical results and interfere with spectral signatures of TP (Leads &Weinstein, 2019). Instead, SEM and energy dispersive X-ray spectroscopy (EDX) or pyrolysis-GC/MS have been suggested as better methods for identifying suspected TP and can provide more accurate characterization of morphological properties in addition to chemical analysis of TP (Sommer et al., 2018; Unice et al., 2012).

## Acknowledgements

This project was conceived as a joint research effort by colleagues from Clemson University (P.van den Hurk), The Citadel (J. Weinstein) and the College of Charleston (B. Beckingham). Assistance with field sampling was provided by Shannon Bley.

Funding was provided by the South Carolina SeaGrant Consortium (R/ER-52), the South Carolina Water Resources Center and Clemson University Creative Inquiry Projects. The authors have no Conflict of Interest to declare.

