## Supplemental Information for "ACCUMULATION OF MICROPLASTIC AND MICRORUBBER PARTICLES IN STORMWATER POND FISH AND INVERTEBRATES"

Table S1. Stormwater Pond Sampling Sites, Mt. Pleasant, SC, USA

| Location | Latitude | Longitude | Type | Receiving Waterbody | Approx. avg daily traffic (# vehicles)[a] |
|---|---|---|---|---|---|
| Patriots Point Links (PP) | 32°47'26.64"N | 79°53'33.56"W | Reference, golf course | Branch of Shem Creek | 225 |
| Whipple Road (WR) | 32°50'05.86"N | 79°50'44.60"W | Residential | Branch of Hobcaw Creek | 11,500 |
| Oak Marsh Drive (OM) | 32°49'53.27"N | 79°51'15.90"W | Residential/Highway | Hobcaw Creek | 49,000 |
| Oyster Point (OP) | 32°49'18.41"N | 79°48'07.29"W | Residential (new) | Branch off Gray Bay | 12,400 |
| Tides Condos (TC) | 32°48'09.03"N | 79°54'02.33"W | Commercial/High Density Residential | Branch of Cooper River | 62,700 |

[a] Approximate average daily traffic (# vehicles) data from South Carolina Department of Transportation (SC DOT) Traffic Counts (2020).

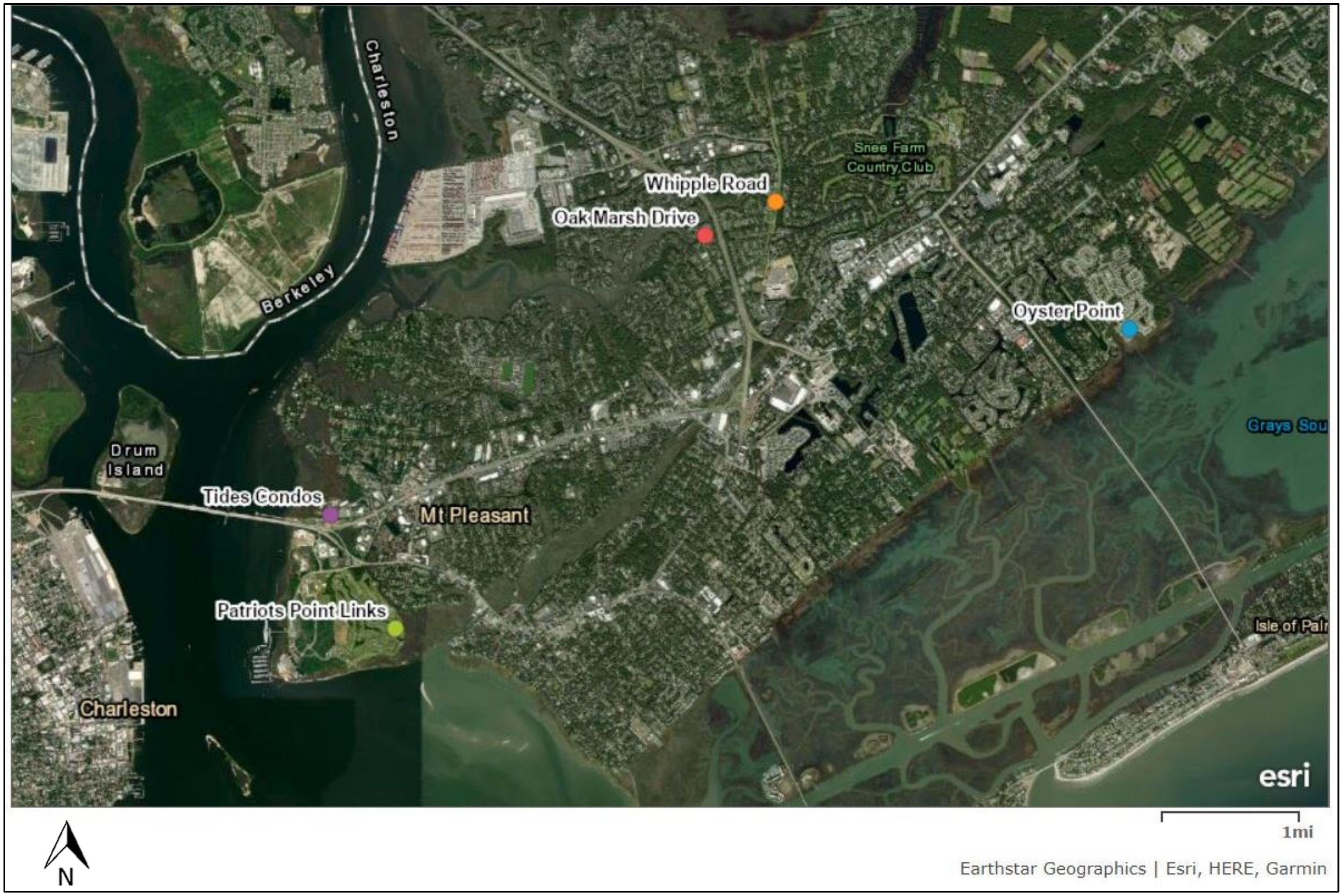


Figure S1: Overview of stormwater pond sampling locations in Mt. Pleasant, SC, USA.

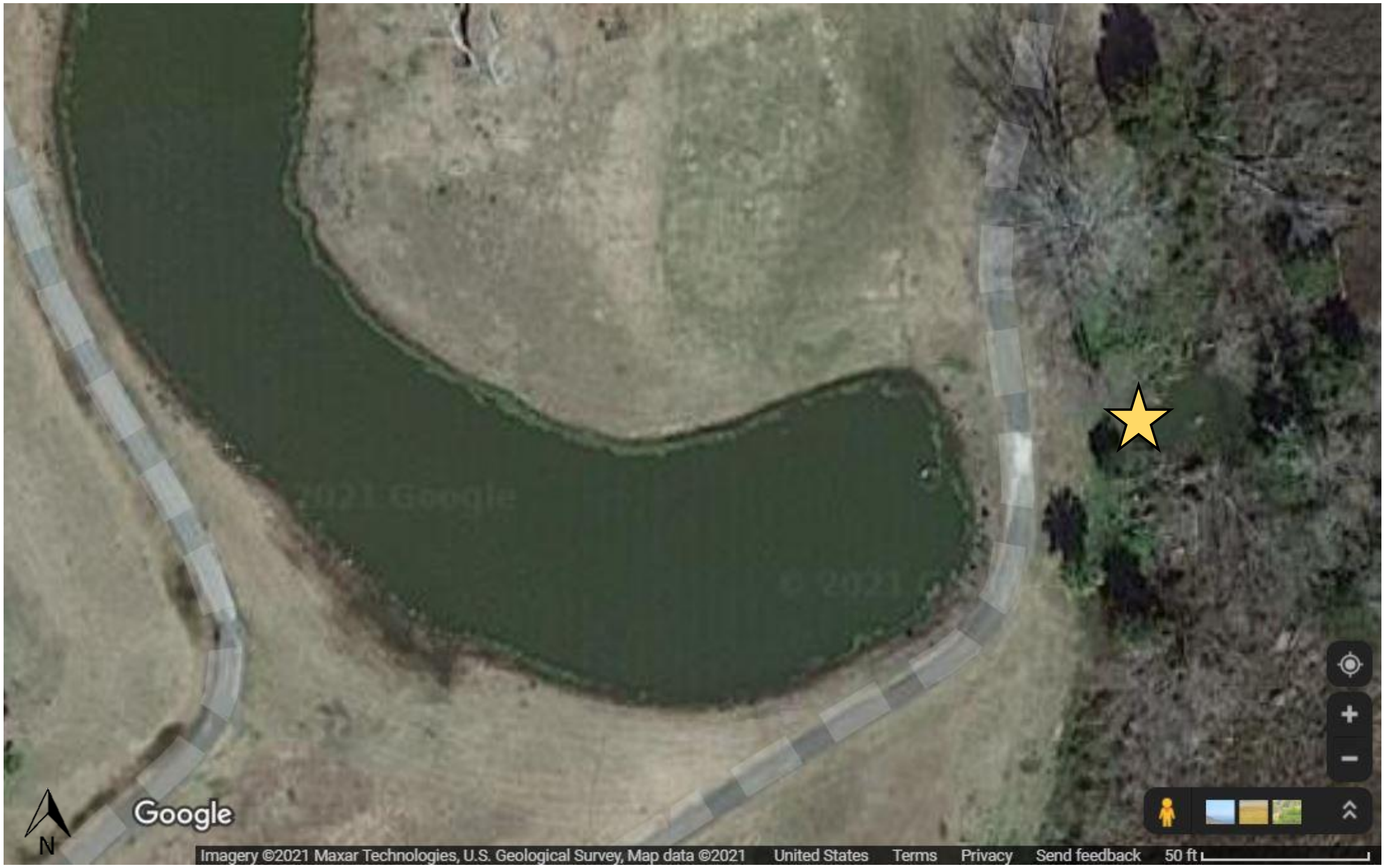


Figure S2: Patriots Point Links (PP) pond and adjacent tidal creek. Star indicates adjacent tidal creek.

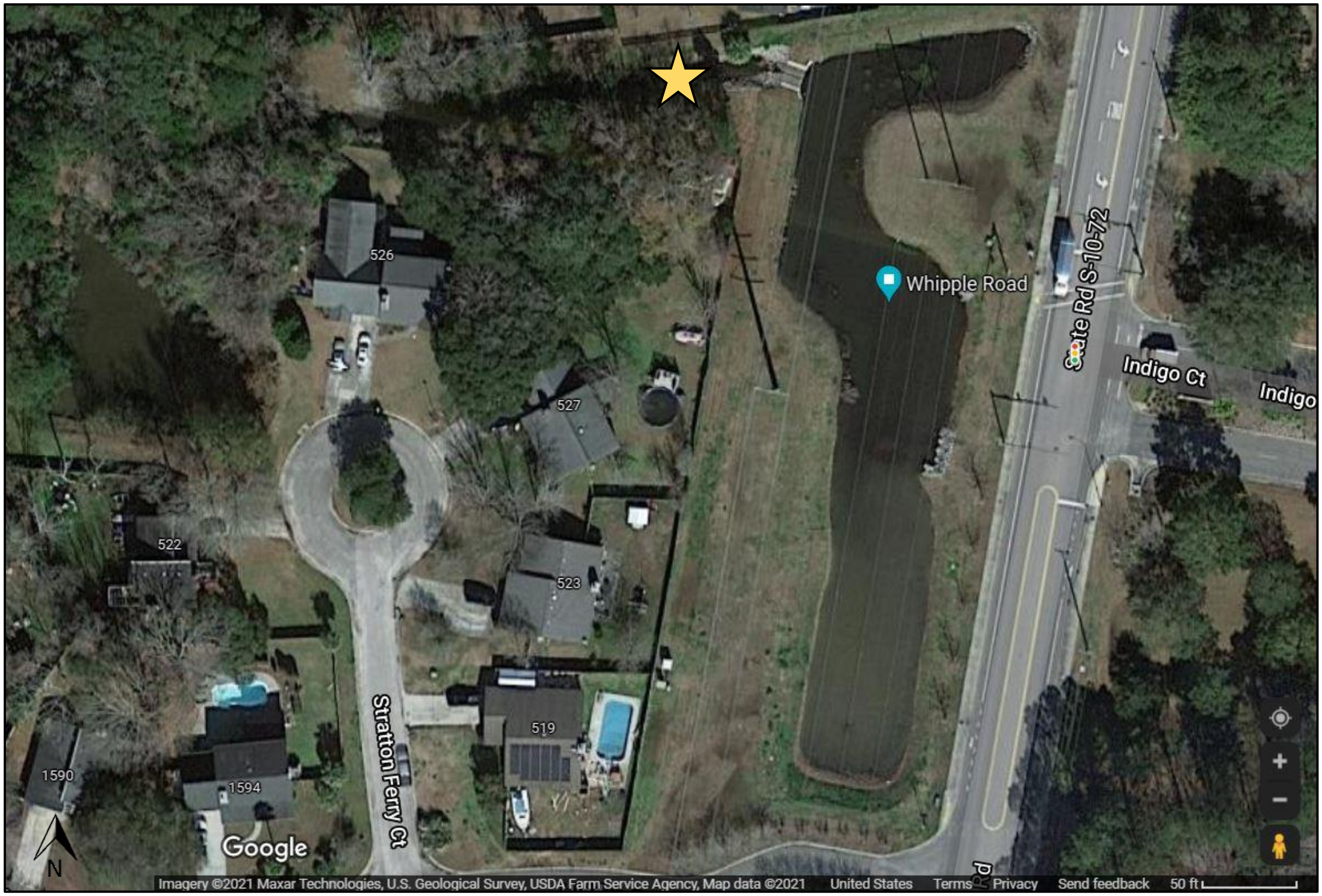


Figure S3: Whipple Road (WR) pond and adjacent tidal creek. Star indicates adjacent tidal creek.

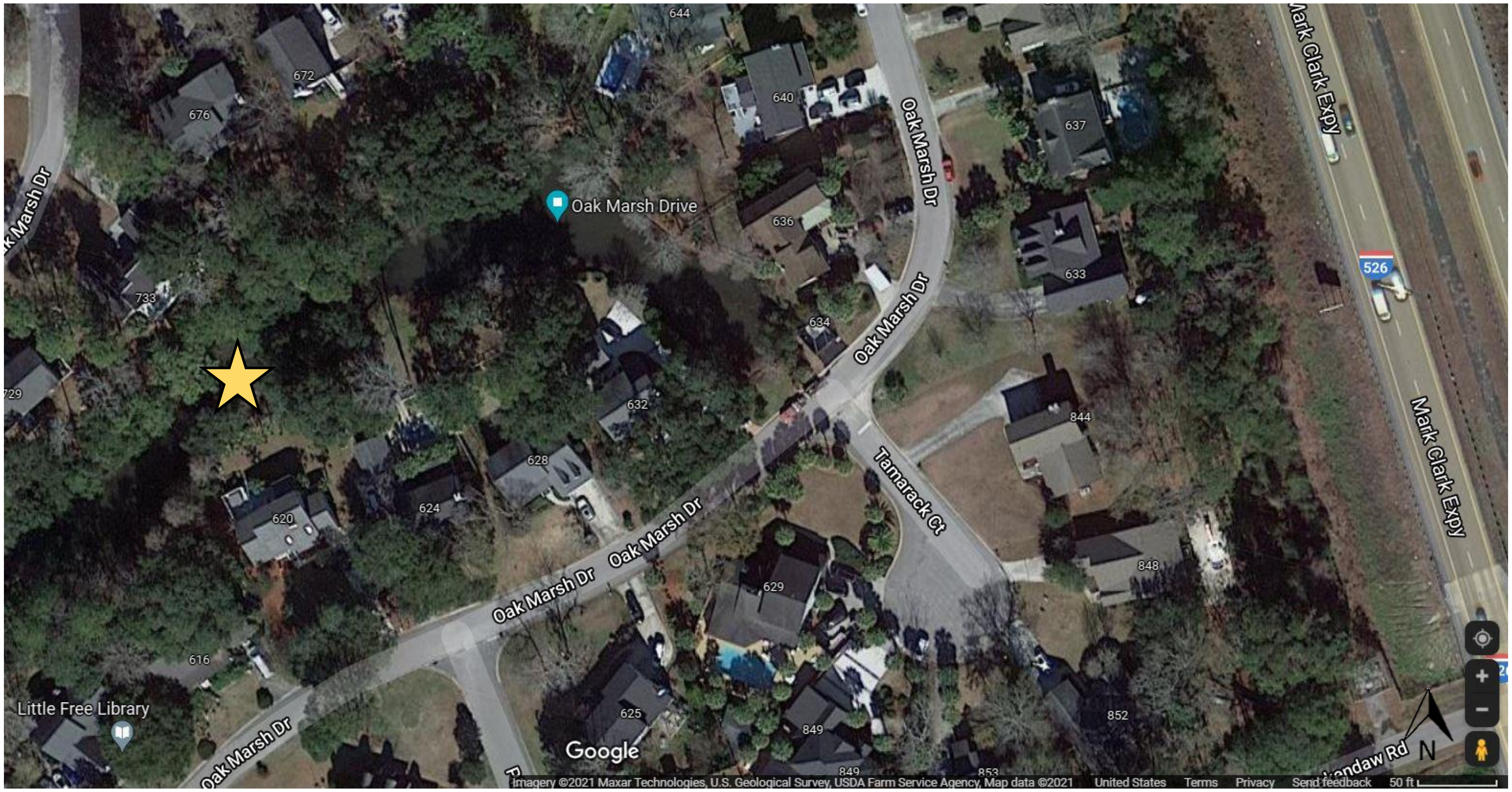


Figure S4: Oak Marsh Drive (OM) pond and adjacent tidal creek. Star indicates adjacent tidal creek.

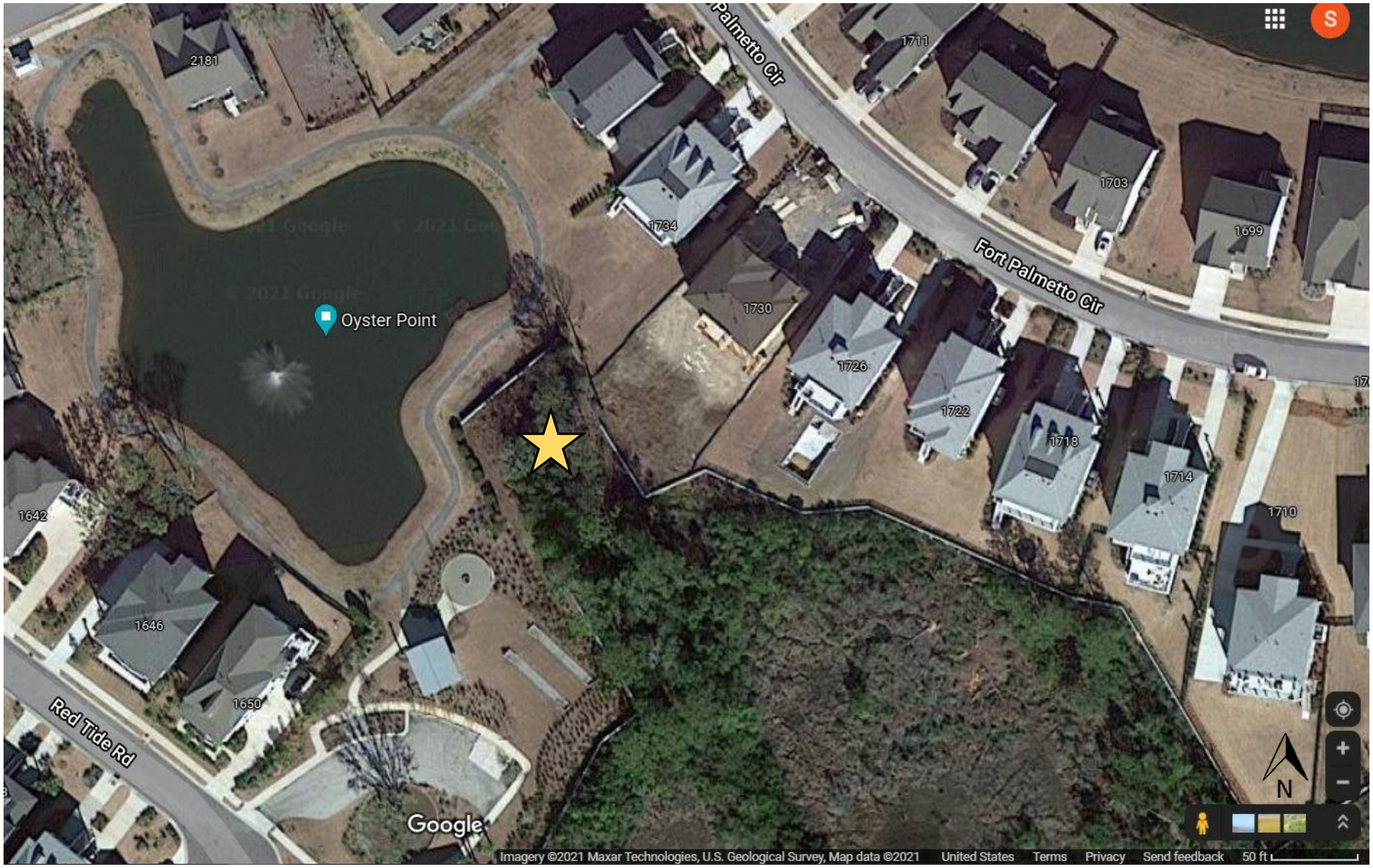


Figure S5: Oyster Point (OP) pond and adjacent tidal creek. Star indicates adjacent tidal creek.

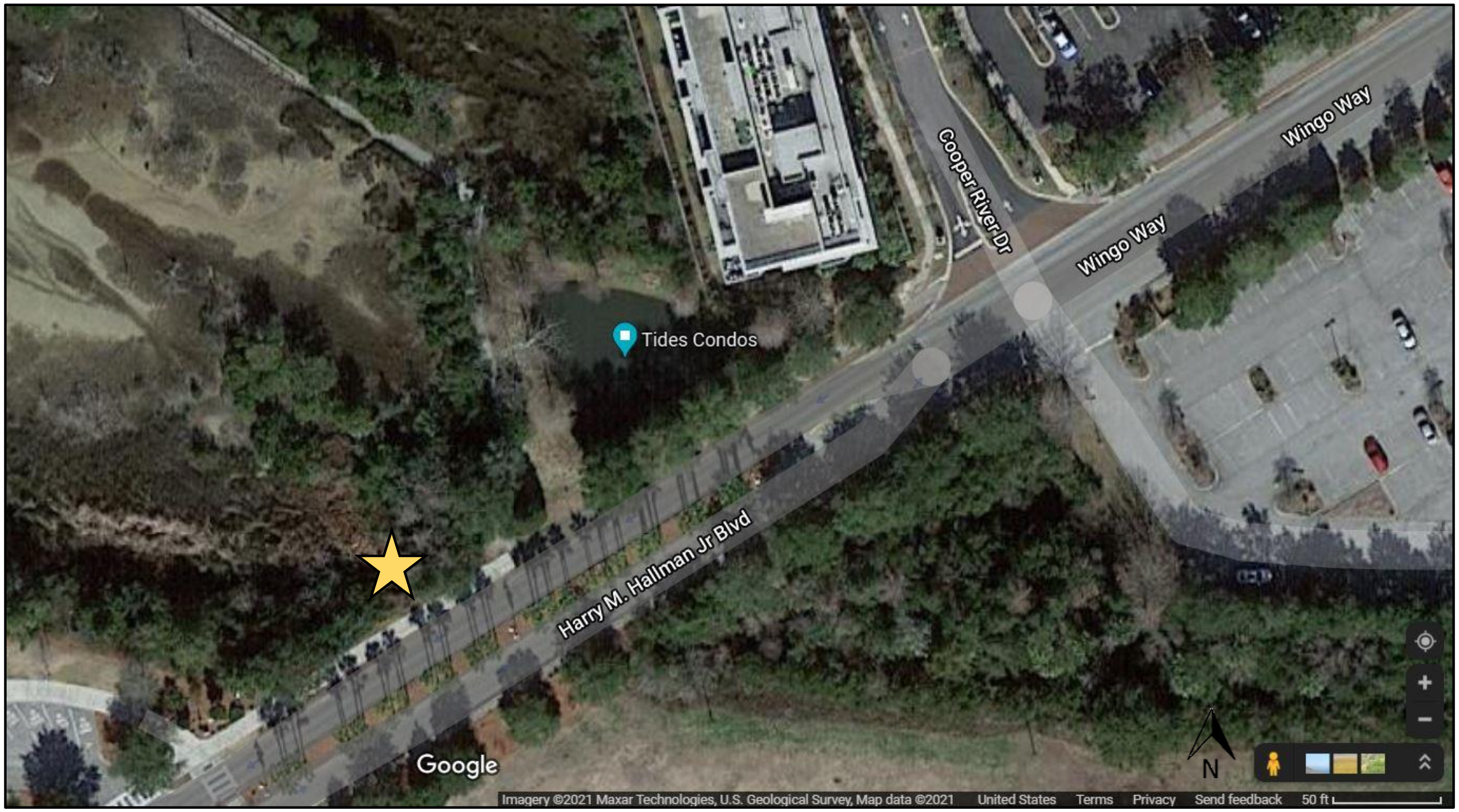


Figure S6: Tides Condos (TC) pond and adjacent tidal creek. Star indicates adjacent tidal creek.

Table S2. Species information for organisms collected from stormwater ponds

Species, common name, family, total number of processed organisms (n) per species, feeding type, feeding habitat, food source. Arithmetic mean values (± standard error (SE)) for wet weight (g) and standard length (mm) for each species are shown. References are listed in main paper reference list.

| Species | Common name | Family | n | Feeding type | Feeding habitat | Main food | Wet weight (g) | Standard length (mm) |
|---|---|---|---|---|---|---|---|---|
| *Micropterus salmoides* | Bass | Centrarchidae | 30 | Omnivorous* | Surface, mid-water, epibenthic | Zooplankton, aquatic insects[a*] | 0.40 ± 0.02 | 31.0 ± 0.58 |
| *Procambarus troglodytes* | Crawdad | Cambaridae | 3 | Omnivorous, detritivores | Benthic, epibenthic | Aquatic vegetation, insect larvae, small fishes, detritus[b] | 11.6 ± 4.0 | n/a |
| *Palaemonetes pugio* | Grass shrimp | Palaemonidae | 25 | Omnivorous, detritivores | Benthic, epibenthic | Detritus, microalgae, mysids, nematodes[c] | 0.25 ± 0.02 | n/a |
| *Mugil cephalus* | Mullet | Mugildae | 21 | Detrivores | Benthic | Detritus, algae, microcrustaceans[d] | 0.79 ± 0.04 | 36.4 ± 0.74 |
| *Poecilia latipinna* | Molly | Poeciliidae | 24 | Planktivorous, Omnivorous | Surface, mid-water, epibenthic | Algae, plant material, aquatic insects[e] | 1.05 ± 0.16 | 37.5 ± 1.32 |
| *Gambusia holbrooki* | Mosquitofish | Poeciliidae | 96 | Omnivorous | Surface | Mosquito larvae, aquatic insects, plant material[f] | 0.73 ± 0.02 | 36.9 ± 0.36 |
| *Fundulus heteroclitus* | Mummichog | Fundulidae | 13 | Omnivorous | Surface, mid-water, epibenthic | Plant material, crustaceans, shrimps, insects, fish[g] | 4.97 ± 2.15 | 65.4 ± 2.15 |
| *Lagodon rhomboides* | Pinfish | Sparidae | 12 | Carnivorous* | Epibenthic | Shrimp, fish eggs, insect larvae, polychaete worms, amphipods[h] | 1.57 ± 0.14 | 45.2 ± 1.07 |
| *Cyprinodon variegatus* | Sheepshead minnow | Cyprinodontidae | 10 | Omnivorous, detritivore | Epibenthic | Shrimp, aquatic insects, detritus[i] | 2.06 ± 0.22 | 41.4 ± 1.59 |

| Continued | | | | | | | | |
|---|---|---|---|---|---|---|---|---|
| **Species** | **Common name** | **Family** | **n** | **Feeding type** | **Feeding habitat** | **Main food** | **Wet weight (g)** | **Standard length (mm)** |
| *Menidia menidia* | Silversides | Atherinopsidae | 40 | Omnivorous | Pelagic | Algae, invertebrates, zooplankton, insects[j] | 1.06 ± 0.14 | 56.15 ± 2.21 |
| *Lepomis spp.* | Sunfish | Centrarchidae | 8 | Omnivorous | Epibenthic | Aquatic insects, zooplankton, plant material[k] | 2.50 ± 0.14 | 54.87 ± 1.26 |

* Applies for juveniles

[a] Miranda & Pugh, 1997; [b] Skelton, 2012;[c] Odum & Heald, 1972; [d] Bester, 2017; [e] Rohde et al., 1994;[f] Rohde et al., 2009;[g] Abraham, 1985; [h] Feinstein, 1975; [i] Robertson & Van Tassell, 2019; [j] Antonucci et al., 2014; [k] Carlander, 1977
